# TOR signaling regulates GPCR levels on the plasma membrane and suppresses the *Saccharomyces cerevisiae* mating pathway

**DOI:** 10.1101/2024.05.09.593412

**Authors:** Nicholas R. Leclerc, Toby M. Dunne, Sudati Shrestha, Cory P. Johnson, Joshua B. Kelley

## Abstract

Target of Rapamycin (TOR) complexes and G-protein-coupled receptors (GPCRs) are crucial signaling hubs that coordinate adaptive responses to environmental inputs. While GPCR-mediated regulation of TOR has been extensively studied, little is known about TOR-mediated regulation of GPCRs. Here, we establish TOR as a regulator of GPCR signaling via its control of receptor endocytosis in the yeast mating system. By pairing fluorescence microscopy with yeast genetic approaches, we identify the machinery that bridges TOR nutrient sensing to GPCR internalization. Furthermore, we find that autophagic machinery is employed during mating to deliver active receptor to the vacuole (lysosome), suppressing the mating pathway. These results demonstrate that TOR regulates the localization and signaling of the yeast mating GPCR in both ligand-dependent and -independent contexts. These pathways are highly conserved, suggesting that TOR-regulation of GPCRs may be a broadly conserved mechanism for integrating competing signals involving metabolic state and external communications.

## INTRODUCTION

The central metabolic regulator Target of Rapamycin (TOR) kinase is conserved across eukaryotes^1^. TOR kinases were first discovered in yeast as Tor1 and Tor2^2,3^. Subsequently, the mammalian mTOR was identified with highly conserved mechanisms of action^4-7^. Tor kinases assemble into two distinct protein complexes, TORC1 and TORC2 (mTORC1 and mTORC2 in mammals)^8^. These complexes regulate metabolism, growth, autophagy, and endocytosis in response to various stressors^9,10^. While mammalian cells harbor one mTOR protein that assembles into either complex, *S. cerevisiae* have two, Tor1 and Tor2^8^. This is likely due to whole genome duplication, a significant evolutionary event that led to the generation of multiple copies of pre-existing proteins^11^. While Tor1 assembles into TORC1, Tor2 may assemble into both TORC1 and TORC2^8,12^.

TORC1 resides in the perivacuolar space^13^ and is primarily responsive to available nutrients, such as amino acids^14^. In nutrient rich environments, TORC1 is active and phosphorylates downstream effectors to enhance anabolism and protein synthesis while repressing catabolism and the cellular recycling process known as autophagy. Bulk (non-selective) autophagy can be induced pharmacologically by rapamycin^15^, a macrolide that forms a complex with Fpr1 (FKBP12 in humans) to directly inhibit TORC1^2,5-7,16^. This results in the formation of double-membraned vesicles, known as autophagosomes, that engulf cytoplasmic cargo as their membranes grow, mature, and close. Autophagosomes ultimately traffic to the vacuole, leading to the destruction of both the vesicle and its internal cargo. This resupplies the cell with free metabolites, prolonging cell survival during nutritional stress^17^.

TORC2 regulates actin polarization, sphingolipid biosynthesis, and endocytosis. TORC2 is localized at the cytoplasmic side of the Plasma Membrane (PM)^18,19^, where it responds to membrane stress^20,21^. Stressors increase the activity of TORC2, promoting sphingolipid synthesis and the cell wall integrity pathway through the AGC kinase Ypk1 (homolog of mammalian SGK1)^22,23^. Aside from its structural and functional differences, TORC2 is dissimilar to TORC1 in two ways: 1) stress upregulates the activity of the complex^23-25^ and 2) it is insensitive to rapamycin^8^. Together, TORC1 and TORC2 each serve as unique signaling nexuses that promote survival of cells facing stressful conditions.

The TOR complexes on their own are crucial sensors of environmental stressors ^9,10,20,26,27^, a function that is necessary to coordinate adaptive responses against unfavorable conditions. G-protein-coupled receptors (GPCRs) are also critical sensors of extracellular cues ^28,29^, many of which organize signaling relays to regulate TOR activity. For instance, several GPCRs that upregulate PKA activity lead to a reduction in mTORC1 activity, such as the β2-adrenergic receptor and the glucagon receptor^30^. GPCR activation may also upregulate mTORC1 activity, such as the amino acid receptors T1R1 and T1R3^31^. However, whether the regulatory effect of GPCRs is positive or negative varies between systems and cell types^32^. While there is a growing wealth of knowledge surrounding GPCR-mediated control of the TOR complexes, little is known about how TOR may regulate GPCR signaling.

GPCRs are a conserved family of eukaryotic membrane proteins that serve as sensors of the extracellular environment^28,29^. GPCRs are functionally unique from other cell surface receptors due to their coupling with a heterotrimeric G-protein (consisting of Gα, Gβ, and Gγ subunits)^33^. This heterotrimeric G-protein facilitates signal transduction from an active receptor to downstream effectors, including adenylate cyclase^34^, phospholipase C^35^, phosphoinositide 3-kinases (PI3K)^36^, and mitogen-activated protein kinases^37^ to induce adaptive responses. In *S. cerevisiae,* mating of haploids is controlled by a GPCR system. These cells secrete pheromones depending on their mating type: MAT**a** cells secrete **a**-factor, and MATα cells secrete α-factor^38-40^. Cells recognize mating pheromone of the opposite type through their mating GPCRs: Ste2 in MAT**a**^41^, and Ste3 in MATα^42^. These two GPCRs initiate identical signaling cascades resulting in polarized growth, cell cycle arrest in G1, and transcription of mating genes. Through this GPCR-mediated response, cells are primed to form a mating projection toward the source of pheromone, fuse, and mate with the nearest available partner of the opposite mating type^38-40^. In the case that multiple cells attempt to mate with the same potential partner, only one can complete the mating process^43^. This poses viability and fitness risks for the cells that respond to pheromone but cannot find a mate. Cells that respond to pheromone for prolonged periods of time are at a greater risk of death^44^. Moreover, cells that continue to try and mate will spend more time, energy, and nutrients forming an obsolete mating projection, giving up the potential to produce more progeny and maintain their biological fitness. Therefore, cells must constrain the activation of their pheromone receptors. Cells have programmed mechanisms to downregulate GPCR signaling following pheromone recognition. This occurs primarily through the 1) inactivation of the G-protein^45,46^ and 2) internalization of the GPCR^47^. These processes desensitize the cell to pheromone at the ligand, receptor, and G-protein level, downregulating the mating pathway and preventing further pheromone signaling.

Endocytosis of the yeast pheromone receptor Ste2 following its activation is well understood. GPCR activation results in C-terminal phosphorylation by Yeast Casein Kinases (Yck1 and Yck2)^48-50^. This phosphorylation allows for the recruitment of α-arrestins (Rod1 and Rog3)^47,51,52^. The receptor tail is then multi-monoubiquitinated^53^, allowing for recruitment of epsin-like proteins (Ent1 and Ent2) that promote assembly of endocytic machinery ^54,55^ and drive internalization of the receptor^56,57^. Following receptor-mediated endocytosis, Ste2 is trafficked to the vacuole (lysosome) for degradation^58,59^, whereas Ste3 localizes at the endosome and is recycled^60^. Both scenarios lead to the desensitization of the mating receptors to pheromone^47,61^.

During the pheromone response, cytoplasmic proteins have been found to accumulate in the vacuole^62^, a hallmark of autophagy despite the lack of nutritional stress. In other studies, proteomic approaches suggested that nitrogen starvation diminishes Ste2 levels^63^. These data led us to hypothesize that TOR inhibition and autophagy machinery may serve to drive Ste2 endocytosis and suppression of the pheromone signaling pathway. Here, we show that TOR signaling regulates the surface presentation of both mating GPCRs in *S. cerevisiae* during TORC1 inhibition, leading to the internalization of the mating receptors. Coincident with this altered receptor localization, we find that TORC1 inhibition dampens pheromone signaling and the ability to mate. Furthermore, we identify key signaling components linking TORC1 to Ste2 internalization, such as TORC2 and its effector Ypk1. We then assess whether TORC effectors are utilized to suppress the mating response following exposure to pheromone. We find that Ypk1 significantly represses the mating pathway. We also show that the central autophagy protein Atg8 aids in the vacuolar targeting of Ste2 to the vacuole, suppressing the mating pathway. Together, we report that TOR complexes and their effectors can regulate GPCRs in both ligand-dependent and ligand-independent contexts.

## RESULTS

### Nutrient availability regulates surface presentation of the mating GPCR Ste2

While both glucose and nitrogen starvation have been reported to drive Ste2 internalization, these two stresses signal through different pathways^63,64^. These pathways could serve as protective mechanisms to prevent yeast from mating in inadequate nutritional conditions via internalization and subsequent destruction of the mating receptors. The signaling that could lead to this is muddled: glucose starvation promotes TORC1 activity while nitrogen starvation suppresses it and TORC1 can upregulate or downregulate endocytosis in different contexts^63-67^. Nitrogen starvation impacts on Ste2 have not been investigated beyond the proteomics screen that found reduced Ste2 levels^63^. Therefore, we first set out to validate the effects of nitrogen starvation signaling on Ste2 internalization. To test this, we examined cells expressing Ste2-mEnvy from a culture grown to saturation. Cells were either grown to the stationary phase of growth (OD600 > ∼1.2-1.5) to create a nutrient-deprived environment or maintained in the nutrient-rich mid-log phase of growth (OD600 < 0.8) and then imaged by fluorescence microscopy (Fig. 1A). Cells in the mid-log phase showed typical receptor localization patterns, Ste2 was present on the PM and inside the vacuole of the cell. However, cells grown to stationary phase showed a 62% reduction in abundance at the PM (Fig. 1B). We tested whether this was a reversible effect by diluting stationary phase cells and supplementing them with fresh nutrients over 4.5 hours. The control group was sustained in mid-log growth by regular media changes. Previously-starved cells showed a complete recovery of PM-associated Ste2 after 1.5 hours in replete media (Fig. 1B). This shows that the available nutrients determines the spatial localization of the mating receptor.

**Fig. 1.**
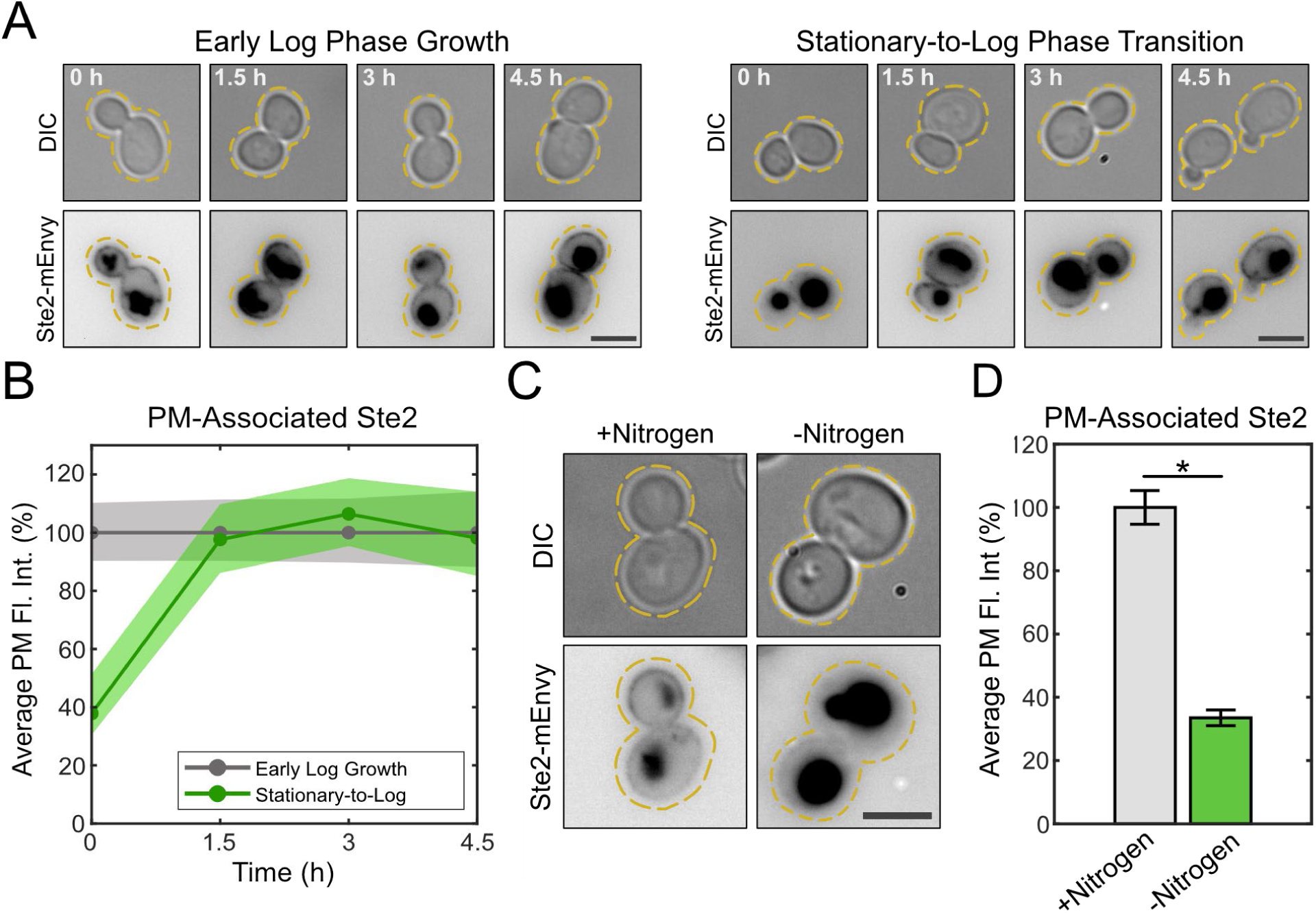
Nutrient availability regulates Ste2 levels at the PM. (**A**) Representative images of cells expressing Ste2-mEnvy maintained in the log phase of growth (left panels) (n > 130 cells) or grown to stationary phase and then diluted in replete media (right panels) (n > 90 cells) for 4.5h. Scale bars represent 4µm. (**B**) Quantification of Ste2-mEnvy mean fluorescent intensity at the plasma membrane every 1.5h either during early log phase growth or growth from stationary phase into replete media. Measurements were normalized to the mean of “Early Log Growth” values at each timepoint to produce percentages. Shaded areas represent 95% bootstrap confidence intervals. (**C**) Representative images of cells expressing Ste2-mEnvy grown in SCD media (+Nitrogen) (n = 252 cells) or low nitrogen SCD media (-Nitrogen) (n = 568 cells) for 6h. Scale bar represents 4µm. (**D**) Quantification of Ste2-mEnvy mean fluorescent intensity at the plasma membrane in cells grown in SCD media (+Nitrogen) or low nitrogen SCD media (-Nitrogen) for 6h (n > 250 cells). Measurements were normalized to the mean of the +Nitrogen group to produce percentages. Error bars represent ±SEM. * = p < 0.001 by *t*-test.

While saturated cultures should be depleted of both nitrogen and glucose, we were specifically interested in TOR signaling and its regulation by nitrogen depletion^14^. Cells were kept at a sub-saturating density (OD600 = 0.2-0.8) in standard Synthetic Complete plus Dextrose (SCD) media and then diluted into low-nitrogen SCD media for 4.5 hours. Cells were constantly maintained in the mid-log phase of growth throughout the experiment and imaged by fluorescence microscopy (Fig. 1C). Cells in nitrogen-limited media showed a 66% reduction of Ste2 abundance at the cell periphery while in the mid-log phase of growth (Fig. 1D). Thus, localization of Ste2 to the PM is specifically sensitive to nitrogen levels in the media.

### Ste2 plasma membrane levels and the mating response is altered by TORC1

Having found that nitrogen availability controls Ste2 levels on the PM, we hypothesize that the nitrogen-sensitive TORC1 may control the cell signaling that leads to changes in receptor localization. Rapamycin specifically represses TORC1 activity, mimicking the effects of nitrogen depletion^2,5-7^. We treated cells with 0.2µM rapamycin in SCD media during mid-log growth for 2 hours, followed by imaging cells expressing Ste2-mEnvy using fluorescence microscopy. (Fig. 2A). Rapamycin treatment led to a 60% reduction in peripheral Ste2 levels (Fig. 2E), showing TORC1 signaling is responsible for starvation-induced internalization of the receptor. These results indicate that TORC1 inhibition promotes Ste2 internalization.

**Fig. 2.**
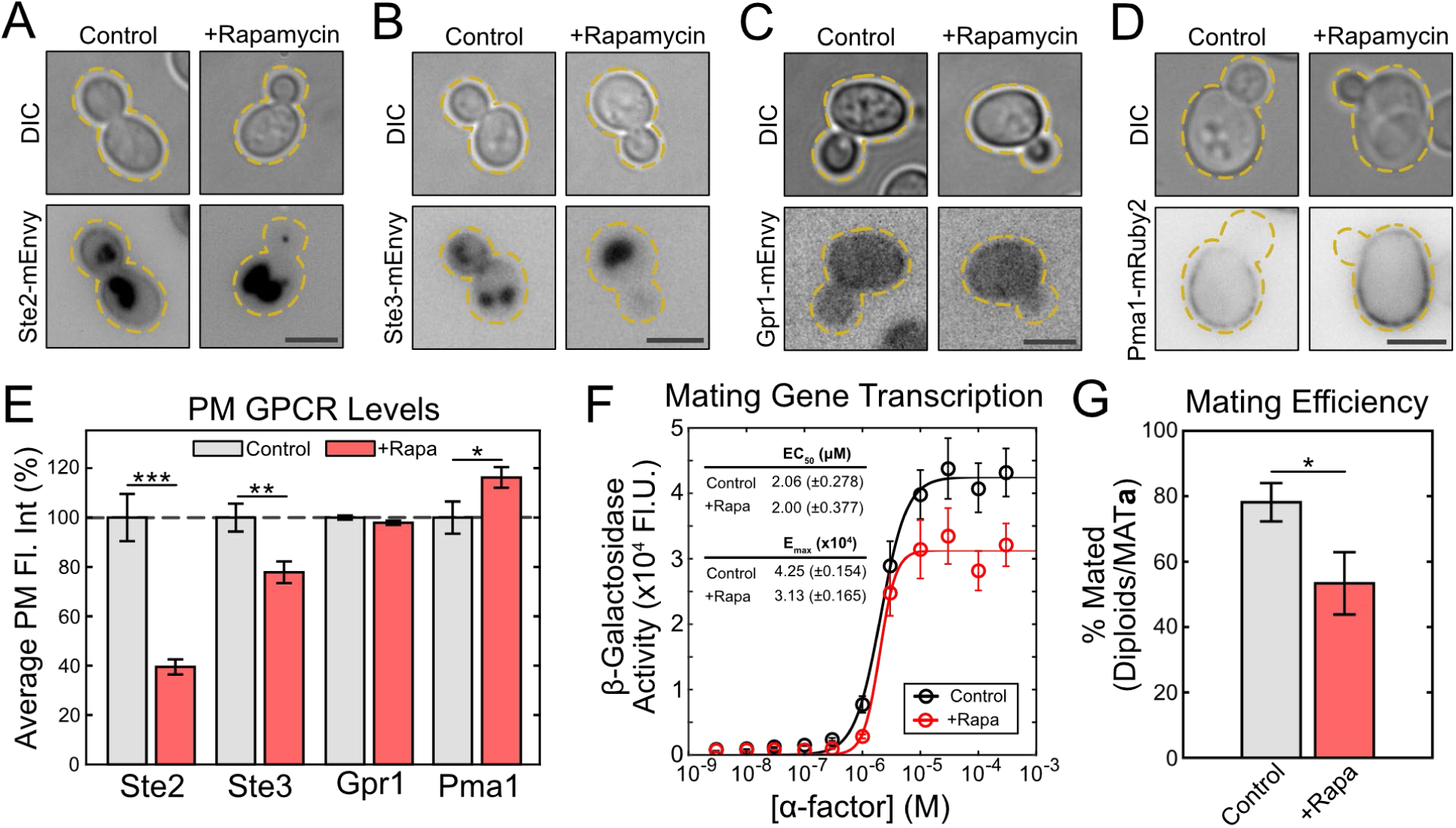
Inhibition of TORC1 reduces mating GPCR levels at the PM and represses the mating pathway. (**A-D**) Representative images of cells expressing (**A**) Ste2-mEnvy, (**B**) Ste3-mEnvy, (**C**) Gpr1-mEnvy, and (**D**) Pma1-mRuby2 treated with or without 0.2µM rapamycin for 2h. Scale bars represent 4µm. (**E**) Quantification of Ste2-mEnvy, Ste3-mEnvy, Gpr1-mEnvy, and Pma1-mRuby2 mean fluorescent intensities at the plasma membrane (-Rap: Ste2 n = 212 cells, Ste3 n = 285 cells, Gpr1 n = 197 cells, Pma1 n = 270 cells. +Rap: Ste2 n = 277 cells, Ste3 n =, Gpr1 n = 214 cells, Pma1 n =). Measurements were normalized to the mean of the untreated groups to produce percentages. Error bars represent ±SEM. * = p < 0.05, ** = p < 0.01, *** = p < 0.001. (**F**) β-galactosidase (pFUS1-LacZ) mating transcription assays of cell cultures pretreated with or without 0.2µM rapamycin for 2h (n > 10 colonies). Error bars represent ±SEM. EC_50_ and E_max_ are reported with ±95% confidence intervals. (**G**) Quantitative mating assays in cells pretreated with or without 0.2µM rapamycin for 2h (n = 5 assays). Error bars represent ±SEM. * = p < 0.05 by *t*-test.

There are three GPCRs in yeast: the mating receptors Ste2 and Ste3, and the sugar receptor Gpr1^68^. To test if TORC1 inhibition impacts other GPCRs, or whether it is selective for Ste2, we generated strains with Ste3 and Gpr1 tagged with mEnvy and treated these cells with 0.2µM rapamycin for 2 hours and then observed their PM levels through fluorescence microscopy (Fig. 2B,C, Supplemental Figure S1). Ste3 levels at the PM decreased by 22%, while Gpr1 abundance remained unchanged (Fig. 2E). To test if other plasma membrane proteins were internalized, we labeled the proton pump Pma1^69^ with mRuby2 and performed the same rapamycin experiment as above (Fig. 2D). Pma1 localization to the PM showed a 16% increase in the presence of rapamycin (Fig 2E). These results show that TORC1 inhibition does not cause indiscriminate endocytosis of integral membrane proteins, as demonstrated by the increase in the abundance of Pma1 and unaltered levels of Gpr1 at the periphery. Further, this TORC1-dependent process is selective among GPCRs, as it specifically downregulates both mating GPCRs Ste2 and Ste3, while not impacting Gpr1.

The reduced surface presentation of Ste2 during TORC1 inhibition led us to hypothesize that rapamycin-treated cells would have a diminished pheromone response. To test this, we measured pheromone-dependent transcription in the presence and absence of rapamycin using a pheromone-responsive β-galactosidase assay (Fig. 2F). In this assay, we use a plasmid with β-galactosidase driven by the pheromone-responsive *FUS1* promoter (pFUS-LacZ)^70^. Therefore, we can measure the dose-response of pheromone-induced transcription based on the activity of the β-galactosidase reporter. We found that rapamycin treatment reduces the maximal transcriptional output (E_max_) of mating genes by 26%. Rapamycin had no effect on the EC_50_. Thus, TORC1 inhibition both removes Ste2 from the membrane and suppresses Ste2-induced transcription.

This lead us to believe that mating should be more difficult to achieve when TORC1 activity is inhibited. To test this, we performed quantitative mating assays^71^ in cells treated with or without rapamycin (Fig. 2G). Under control conditions, 78% of untreated cells can successfully mate, consistent with previously reported levels^72^. In contrast, only 53% of cells treated with rapamycin successfully mated. Altogether, this supports the idea that TORC1 inhibition serves to diminish the mating response, in part through the internalization of mating receptors.

### TORC1-dependent endocytosis occurs through Clathrin-Mediated Endocytosis (CME) and TORC2 signaling

We set out to determine the components in the pathway linking TORC1 inhibition to loss of Ste2 from the PM. To accomplish this, we constructed fifteen strains with deletions of genes known to be involved in TORC signaling, Ste2 endocytosis, and clathrin-mediated endocytosis (CME) (Fig. 3A). These cells were treated with 0.2µM rapamycin for 2 hours, and peripheral Ste2 levels were measured using fluorescence microscopy. Each deletion mutant was then grouped into one of three categories based on whether they were statistically equivalent (by ANOVA followed by multiple comparison) to the control, the rapamycin treatment, or neither when treated with rapamycin. The three groups are: 1) no rescue (not different from WT +rapamycin), 2) partial rescue, and 3) complete rescue (Fig. 3B, Supplemental Figures S1 and S2).

**Fig. 3.**
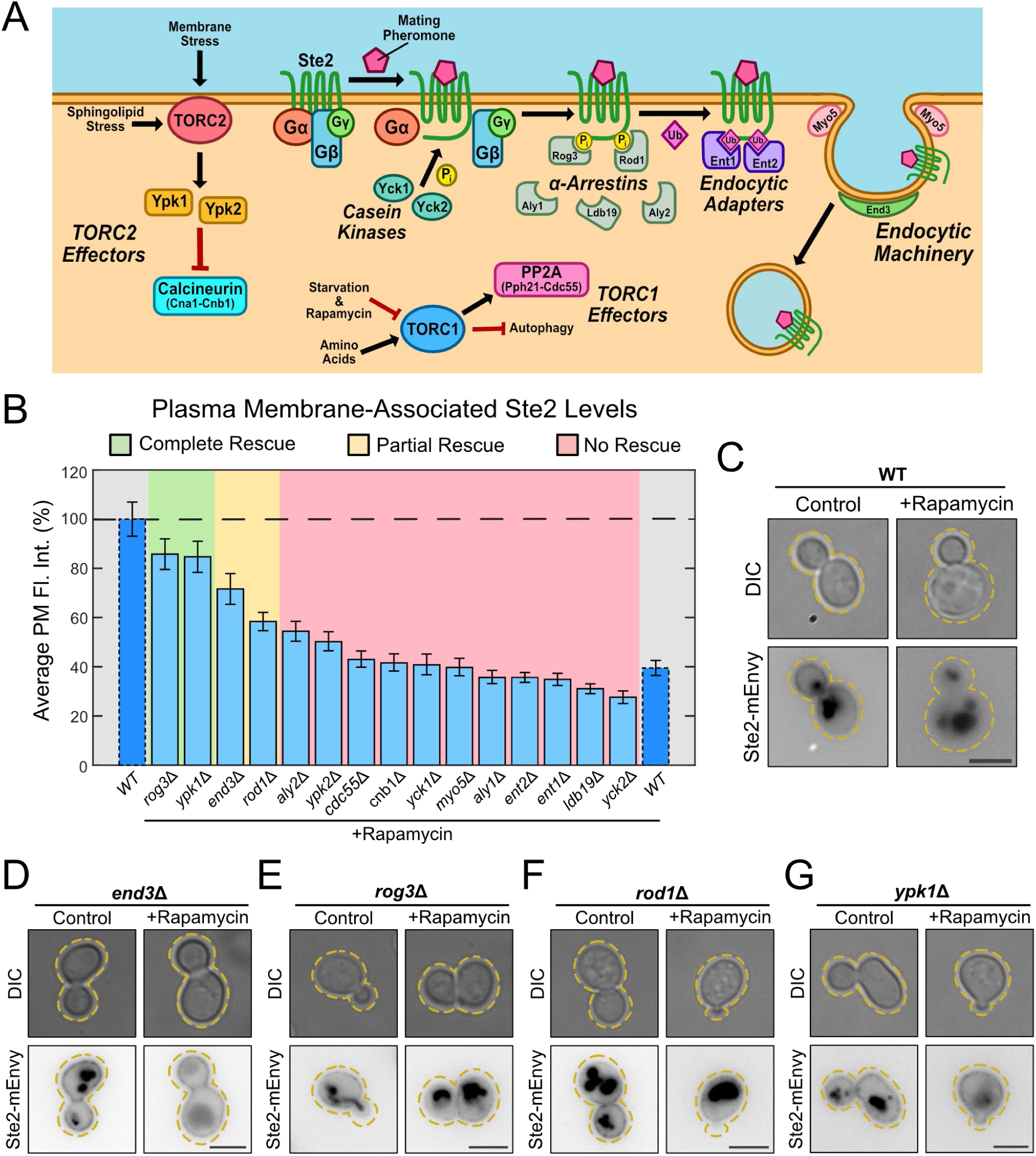
TORC1 inhibition leads to CME of Ste2 through α-arrestins and Ypk1. (**A**) To identify candidates involved in TORC1-mediated endocytosis, we deleted multiple genes that are involved in TORC signaling, receptor endocytosis, and CME. Casein kinases ^48-50^, α-arrestins ^47,51,52,112^, and epsin-like proteins ^56,57^ are all involved in priming the pheromone receptors for CME during mating. CME utilizes motors ^113^ and cytoskeletal organizers ^73,74^ to form then endocytic pit that matures into an endosome. During starvation, TORC1 activity is reduced, downregulating PP2A activity ^114^. TORC2 activates Ypk1 and Ypk2 ^20,22,23^, which repress major cell regulators such as Calcineurin ^115^. (**B**) Quantification of mean Ste2-mEnvy on the plasma membrane in the indicated strain, with or without 0.2µM rapamycin for 2h (n > 170 cells). Measurements were normalized to the mean of the untreated groups (not shown) to produce percentages. One-Way ANOVA was used to analyze variance between treated and untreated mutant and WT cells. Tukey’s honest significance test was used as a multiple comparison test to compare means between groups. Based on ANOVA results, genetic mutants treated with rapamycin were grouped into one of three categories: No Rescue (red), Partial Rescue (yellow), and Complete Rescue (green). Mutants in the No Rescue category were only significantly different (p < 0.05) from untreated WT cells. Mutants in the Partial Rescue category were significantly different from both treated and untreated WT cells. Mutants in the Complete Rescue category were only significantly different from rapamycin-treated WT cells. Error bars represent ±SEM. Bars with dashed outlines show data reported in Figure 2E. (**C-G**) Representative images of (**C**) WT, (**D)** *end*3Δ, (**E**) *rog*Δ, (**F**) *rod*1Δ, and (**G**) *ypk*1Δ cells expressing Ste2-mEnvy treated with or without 0.2µM rapamycin for 2h. Scale bars represent 4µm.

We found that deleting End3 partially blocks the effect of rapamycin on Ste2 localization on the PM (Fig. 3B, D). End3 is an adaptor protein that facilitates the internalization step in CME. Cells that lack End3 show defects in endocytosis and have been reported to reduce internalization of Ste2 by ∼60-80%^73,74^. CME in these cells is defective, but not entirely absent. We also saw that deletion of Rod1 partially blocked the effect of rapamycin on Ste2 localization (Fig 3B,F). In contrast, deleting the α-arrestin Rog3 completely blocked rapamycin-induced internalization of Ste2 (Fig 3B, E). The α-arrestins Rod1 and Rog3 are paralogs that facilitate CME and desensitization of active Ste2 in response to pheromone^47,51^. The impact of deleting Rod1, Rog3, or End3 all indicate that CME plays a role in the internalization of Ste2 in response to TORC1 inhibition.

Rod1 and Rog3 are known to associate with the phosphorylated C-terminus of active Ste2^47,51^, suggesting that C-terminal modification of Ste2 is required for this internalization event. To test this, we replaced the C-terminus of Ste2 with EGFP (*ste2^T326^-EGFP*) in cells and then treated with rapamycin as described above (Fig. 4A). Ste2^T326^ levels at the PM did not change when treated with rapamycin (Fig. 4B, Supplemental Figure S1), confirming that the internalization of the receptor is reliant on the C-terminal tail.

**Fig. 4.**
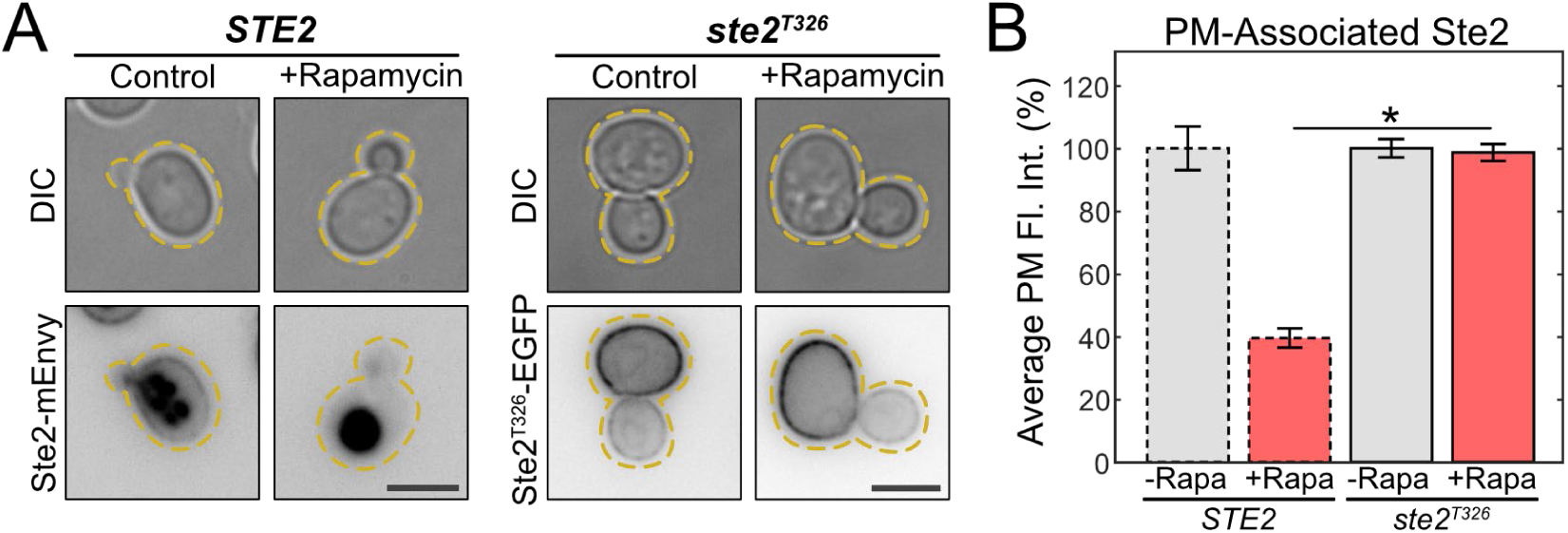
The C-terminus of Ste2 is required for TORC1-mediated endocytosis. (**A**) Representative images of cells expressing Ste2-mEnvy and Ste2^T326^-EGFP treated with (n = 184 cells) and without (n = 132 cells) 0.2µM rapamycin for 2h. Scale bar represents 4µm. (**B**) Quantification of Ste2-mEnvy and Ste2^T326^-EGFP mean fluorescent intensities at the plasma membrane. Measurements were normalized to the mean of the untreated groups to produce percentages. One-Way ANOVA paired with Tukey’s honest significance test was used to compare means between groups. Error bars represent ±SEM. * = p < 0.001. Bars with dashed outlines show data reported in Figure 2E. Scale bars represent 4µm.

Finally, we found that deleting Ypk1 completely blocks the effect of rapamycin on Ste2 PM levels (Fig 3B,G). This was surprising because Ypk1 is a kinase that acts downstream of Pkh1, Pkh2, and TORC2. Pkh1 and Pkh2 are kinases that phosphorylate Ypk1 in its activation loop, a process that is required for its activity^75-77^. Therefore, we deleted Pkh1 and Pkh2 in cells expressing Ste2-mEnvy and treated them with rapamycin to see if upstream components of Ypk1 also rescue Ste2 abundance at the plasma membrane (Fig. 5A,C). We found that both deletions partially rescued Ste2 abundance at the plasma membrane (Fig. 5B,D, Supplemental Figure S1).

**Fig. 5.**
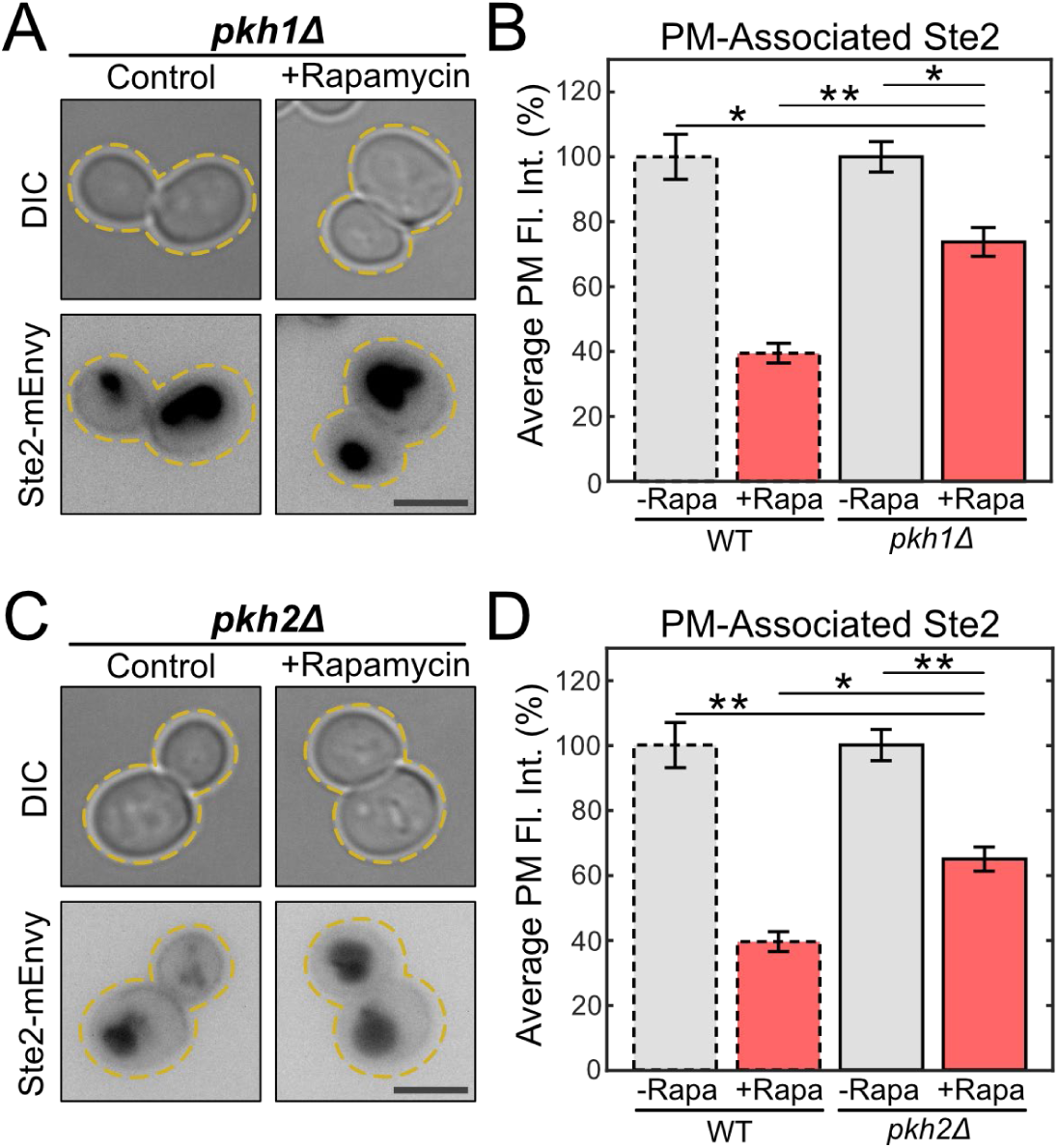
The Ypk1-activators Pkh1 and Pkh2 contribute to TORC1-mediated Ste2 endocytosis. (**A**) Representative images of *pkh1Δ* cells expressing Ste2-mEnvy treated with (n = 186 cells) or without (n = 195 cells) 0.2µM rapamycin for 2h. (**B**) Quantification of Ste2-mEnvy mean fluorescent intensities at the plasma membrane. Measurements were normalized to the mean of the untreated groups to produce percentages. One-Way ANOVA paired with Tukey’s honest significance test was used to compare means between groups. (**C**) Representative images of *pkh2Δ* cells expressing Ste2-mEnvy treated with (n = 135 cells) or without (n = 162 cells) 0.2µM rapamycin for 2h. (**D**) Quantification of Ste2-mEnvy mean fluorescent intensities at the plasma membrane. Measurements were normalized to the mean of the untreated groups to produce percentages. One-Way ANOVA paired with Tukey’s honest significance test was used to compare means between groups. All error bars represent ±SEM. * = p < 0.01, ** = p < 0.001. All bars with dashed outlines show data reported in Figure 2E. Scale bars represent 4µm.

Pkh1 and Pkh2 are functionally redundant^75^, therefore it is unsurprising that the rescue seen in either strain is not complete. However, their individual absences demonstrate a sufficient effect to indicate their necessity for Ste2 internalization during TORC1 inhibition, most likely through Ypk1 activation.

Ypk1 is upregulated by TORC2 in response to stress^24-26,77^. We next tested whether TORC2 signaling was required for Ste2 internalization in response to TORC1 inhibition. The core protein of TORC2, Tor2, is essential, and TORC2 is not normally sensitive to rapamycin^8^. Therefore, we could not delete Tor2 or inhibit it as in our other experiments. Instead, we made TORC2 sensitive to rapamycin by truncating the TORC2 component Avo3 ^78,79^. We reasoned that if TORC2 is responsible for Ste2 internalization upon inhibition of TORC1, then nitrogen starvation would still drive Ste2 internalization in the Avo3 truncation mutant, as TORC2 function should be maintained. However, rapamycin treatment would now inhibit signaling through both TORC1 and TORC2, and Ste2 would remain on the PM during rapamycin treatment. We performed both nitrogen starvation and rapamycin treatment as above. As expected, nitrogen starvation was still able to drive Ste2 internalization in the *avo3^T1273^* strain (Fig 6A,B). However, when *avo3^T1273^*cells were treated with Rapamycin, Ste2 did not display normal rapamycin-induced internalization (Fig. 6C,D, Supplemental Figure S1). These results together confirm that inhibition of TORC1 promotes Ste2 internalization through TORC2 signaling.

**Fig. 6.**
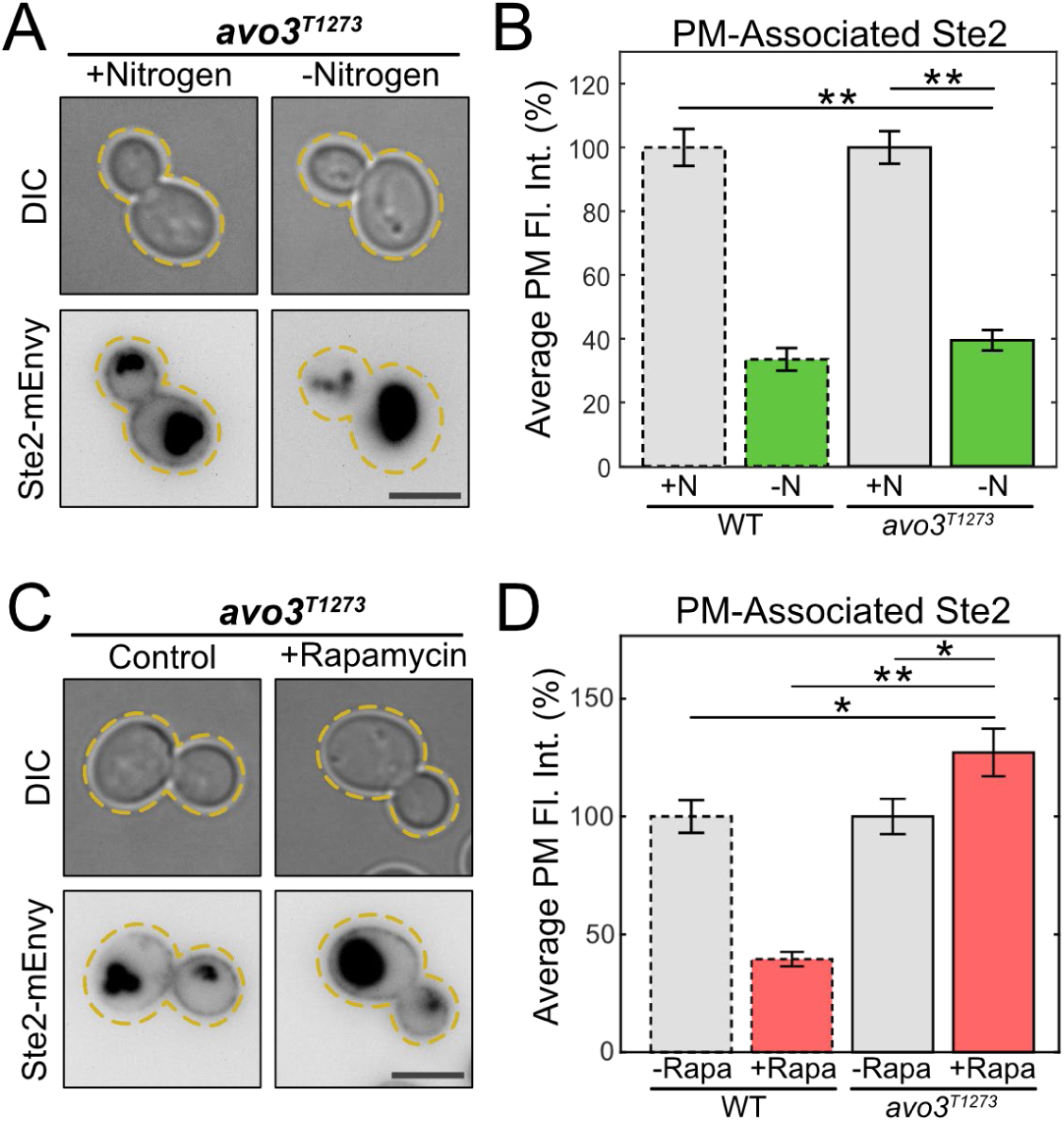
TORC2 activity directs TORC1-mediated Ste2 endocytosis. (**A**) Representative images of cells expressing Ste2-mEnvy and avo3^T1273^ treated with (n = 233 cells) and without (n = 139 cells) 0.2µM rapamycin for 2h. (**B**) Quantification of Ste2-mEnvy mean fluorescent intensities at the plasma membrane. Measurements were normalized to the mean of the untreated groups to produce percentages. One-Way ANOVA paired with Tukey’s honest significance test was used to compare means between groups. (**C**) Representative images of cells expressing Ste2-mEnvy and avo3^T1273^ grown in SCD media (+N) (n = 141 cells) or low nitrogen SCD media (-N) (n = 109 cells) for 6h. (**D**) Quantification of Ste2-mEnvy mean fluorescent intensities at the plasma membrane. Measurements were normalized to the mean of the untreated groups to produce percentages. One-Way ANOVA paired with Tukey’s honest significance test was used to compare means between groups. All error bars represent ±SEM. * = p < 0.05, ** = p < 0.001. All bars with dashed outlines show data reported in Figure 2E. Scale bars represent 4µm.

### Ypk1 suppresses mating transcription in response to pheromone

We have found that Ypk1 is required for rapamycin-induced Ste2 endocytosis. We hypothesize that Ypk1 may be responsible for a concomitant reduction in mating transcription during the pheromone response. To test this, we performed a β-galactosidase assay as described above to measure the transcriptional output of cells lacking Ypk1 (Fig. 7B). These cells show a remarkable 433% increase in transcription of mating genes and a 39% decrease in EC_50_. This indicates that Ypk1 is a negative regulator of the pheromone pathway in the absence of rapamycin or nutritional stress.

**Fig. 7.**
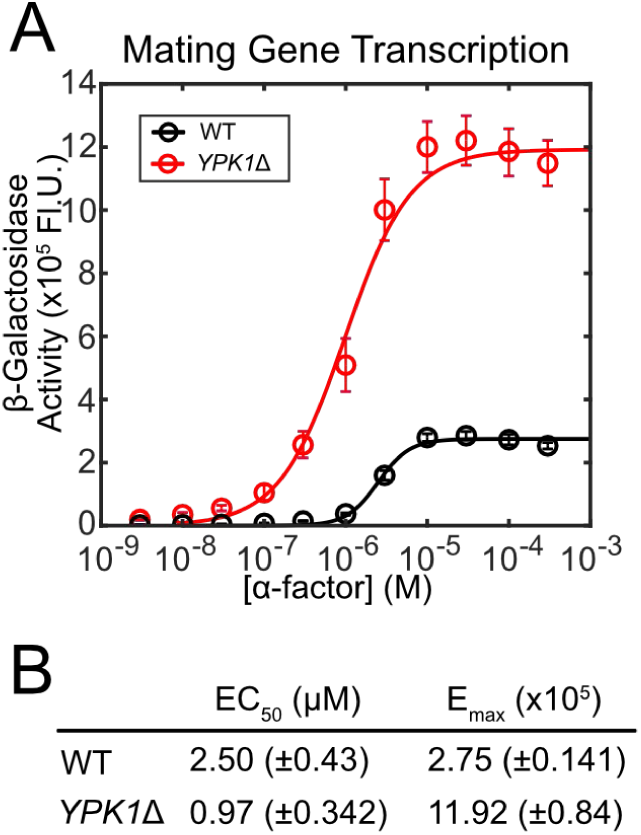
Ypk1 dampens mating pathway output. (**A**) β-galactosidase (pFUS1-LacZ) mating transcription assays of WT (n = 9 colonies) and *ypk1*Δ (n = 9 colonies) cultures. Error bars represent ±SEM. (**B**) Table of EC_50_ and E_max_ responses to α-factor. EC_50_ and E_max_ are reported with ±95% confidence intervals.

### Pheromone-induced autophagy represses mating transcription

Ligand binding to Ste2 drives its internalization^59^ and vacuolar targeting^80,81^. The PI3K Vps34, a protein required for autophagy and vacuolar targeting^82-84^, is also activated during mating^62^. It is possible that autophagy may be activated during mating to help direct Ste2 to the vacuole following its activation and internalization. Here, we have already shown that TORC inhibition, a process which also promotes autophagy, leads to internalization of pheromone receptors in the absence of ligand. We therefore hypothesized that autophagy components could lead to vacuolar targeting of Ste2 during mating. We tested this by deleting the core autophagy protein Atg8 from cells expressing Ste2-mEnvy and Vph1-Tomato and treated them with 10µM pheromone for 2 hours (Fig. 8B). The most notable effect we observed was cells lacking Atg8 appeared to frequently have empty vacuoles. To quantify this, linescans were drawn through the center of the vacuole to measure the levels of Ste2-mEnvy at the vacuolar membrane, labeled by Vph1-Tomato, and the lumen (Fig. 8A). We found that deleting Atg8 led to a 4-fold increase in cells with vacuoles lacking significant Ste2 accumulation (Fig. 8C). This would suggest that during mating, autophagy machinery contributes to the vacuolar targeting of Ste2.

**Fig. 8.**
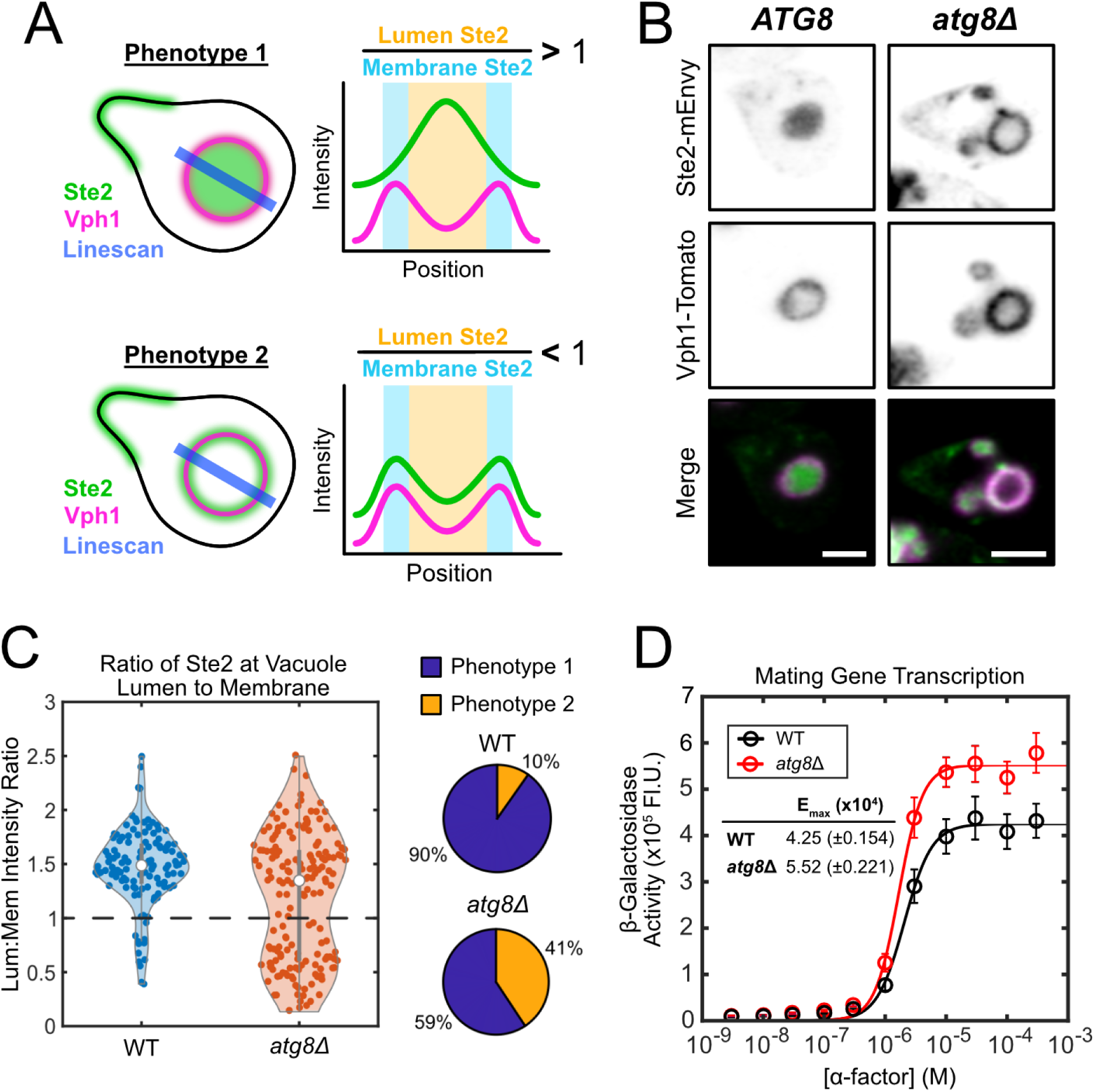
Pheromone-induced autophagy aids in vacuolar targeting of Ste2 and dampens the mating pathway. (**A**) Scheme representing the analysis of Ste2-mEnvy at the vacuolar lumen (Vph1-Tomato) and membrane. We have found that mating cells show two primary phenotypes regarding Ste2 localization to the vacuole. Phenotype 1 shows a cell with a vacuole filled with Ste2-mEnvy, whereas Phenotype 2 shows a empty vacuole, with more receptor at the vacuolar periphery. Linescans were measured through the center of the largest vacuolar body. The mean fluorescent intensity of Ste2-mEnvy was recorded at the peaks of Vph1-Tomato florescence, thereby reporting Ste2 levels at the vacuolar membrane. The mean fluorescent intensity of Ste2-mEnvy between these peaks was reported as Ste2 levels in the vacuolar lumen. The lumen:membrane ratio of Ste2 would then represent the phenotype each cell presents. Cells with Lum:Mem > 1 presented as Phenotype 1 (full), whereas cells with a ratio < 1 presented at Phenotype 2 (empty). (**B**) Representative images of *ATG8* and *atg8Δ* cells expressing Ste2-mEnvy and Vph1-Tomato treated with 10µM α-factor. Scale bar represents 2µm. (**C**) Violin plots and pie charts representing the distribution of WT (n = 123 cells) and *atg8Δ* (n = 172 cells) cells presenting either Phenotype 1 or Phenotype 2 when treated with 10µM α-factor. (**D**) β-galactosidase (pFUS1-LacZ) mating transcription assays of WT (n = 15 colonies) and *atg8*Δ cultures (n = 9 colonies). Error bars represent ±SEM. E_max_ is reported with ±95% confidence intervals.

**Fig. 9.**
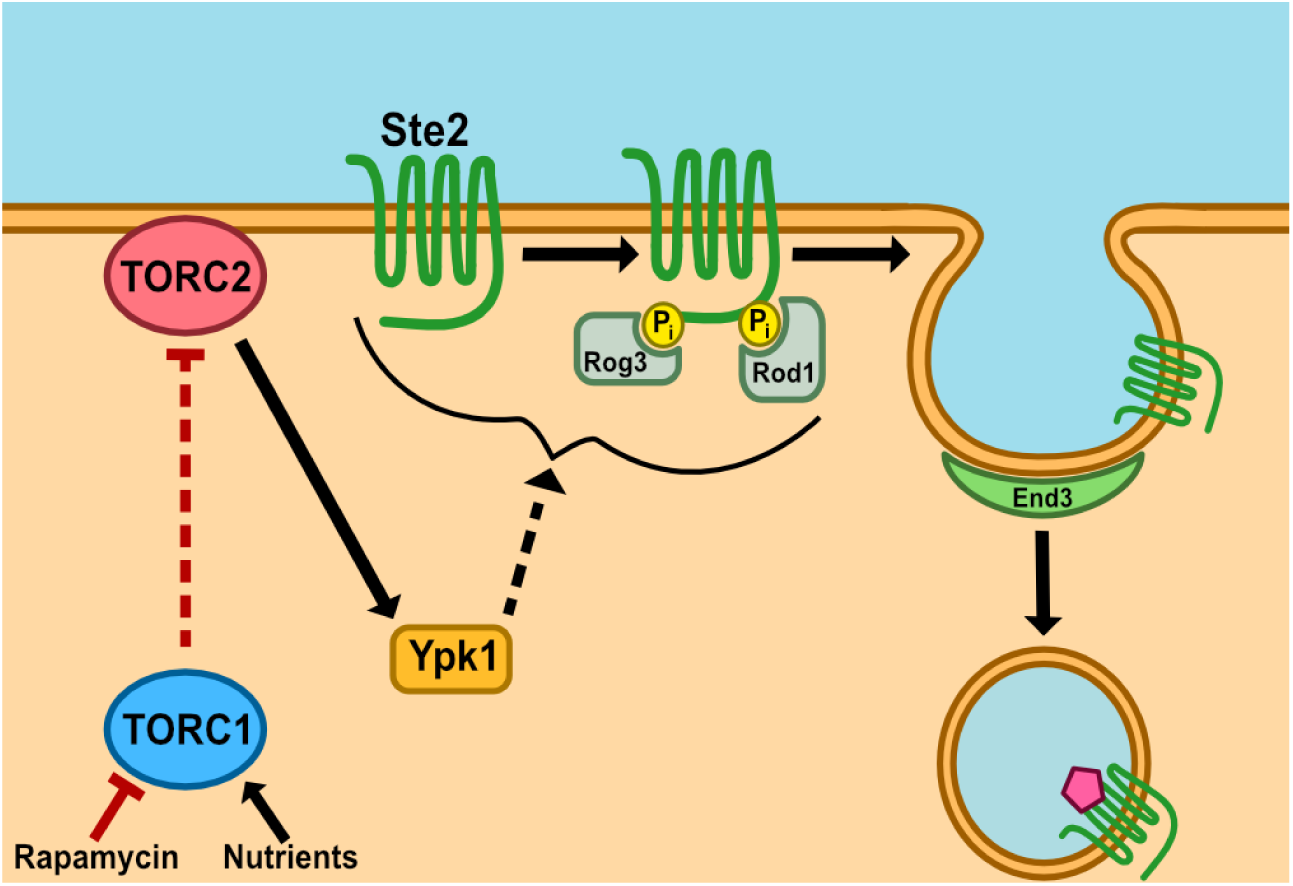
Proposed mechanism of TORC1-directed endocytosis. In nutrient-deprived conditions, TORC1 is repressed, leading to the TORC2-directed activation of Ypk1. Active Ypk1 facilitates α-arrestin-directed CME of Ste2, which ultimately traffics to the vacuole.

Localization of Ste2 to the lumen of the vacuole would remove its cytoplasmic domain from the cytosol and stop any potential endomembrane signaling. We hypothesized that ATG8 and the autophagy machinery may therefore reduce pheromone signaling. To test this, we measured the pheromone dose-response curve with the pheromone-responsive FUS1 β-galactosidase assay described above to measure mating gene transcription in *atg8*Δ and wild-type cells (Fig. 8D). Cells lacking Atg8 showed a 30% increase in transcriptional output compared to wild-type cells, without shifting the EC_50._ Therefore, autophagy during the pheromone response suppresses the total pathway output of pheromone-induced transcription.

## DISCUSSION

Here we describe both ligand-dependent and ligand-independent regulation of the GPCR, Ste2, by TOR complexes and their effectors. We have shown that nutrient limitation leads to the a decrease in the PM levels of mating pheromone receptors Ste2 and Ste3 in *S. cerevisiae*. We found that changes in TOR signaling are responsible for reduced Ste2 abundance on the PM, as rapamycin treatment phenocopies nitrogen limitation. Receptor internalization is concomitant with decreased signaling through the pheromone response pathway as well as a reduction in the ability of the yeast to mate. Through analysis of 15 different candidate genes by deletion analysis and additional truncations of Avo3 and Ste2 itself, we determined that the signaling pathway from TORC1 inhibition to loss of Ste2 from the PM was dependent upon TORC2-Ypk1 signaling and α-arrestin recruitment to the receptor, followed by End3-dependent clathrin-mediated endocytosis. Using pheromone-responsive transcriptional assays, we found that TORC1 signaling promotes pheromone signaling and that Ypk1 signaling suppresses it. Thus, TOR signaling regulates Ste2 PM abundance, mating pathway signaling, and mating itself. As TORC1 inhibition reduces mating, we propose that this signaling pathway serves as a mechanism to suppress mating during times of nutritional stress, when the metabolic requirements of mating may represent a risk to cell survival and fitness.

The link between TOR signaling, autophagy, and endomembrane transport led us to test whether autophagy machinery contributes to receptor trafficking to the vacuole during the pheromone response. We found that the central autophagy protein, Atg8, promotes vacuolar targeting of Ste2 and suppress pheromone-dependent transcription. This clearance of a GPCR from the cytosol through a mechanism that depends on autophagy machinery, while not experiencing nutritional stress, represents a novel mode of negative regulation of GPCR signaling.

### Why suppress mating with TOR?

Cells may reduce their mating receptor levels in response to starvation to avoid mating while hungry. There is an increased risk of cell death while responding to pheromone^44^. If cells are stressed due to a lack of carbon sources and metabolites, mating may exacerbate this stress by committing significant energy and resources toward forming a mating projection. In this situation, it would be advantageous to prevent the mating response. Since Ste2 and Ste3 are at the start of the pathway, they are prime candidates for regulation. In this context, the observation that both Ste2 and Ste3 are removed from the PM suggests that the cell utilizes TOR signaling pathways to inhibit mating when nutrients are scarce.

### TOR-dependent endocytosis vs Ligand-Induced Endocytosis

We have found that TORC1-induced endocytosis of Ste2 is a distinct process from pheromone-induced receptor internalization. During the pheromone response, Ste2 internalization is mediated primarily by three sets of proteins that interact with the receptor C-terminus: 1) the yeast casein kinases Yck1 and Yck2^48-50^, 2) the α-arrestins Rog3 and Rod1^47,51,52^, and 3) the epsin-like proteins Ent1 and Ent2^56,57^. Starvation-induced endocytosis aligns with receptor mediated internalization in that 1) the receptor C-terminus is a required component, 2) it is mediated by α-arrestins, and 3) it is clathrin-mediated as shown by our deletion screen hits (End3, Rog3, and Rod1) and ste2^T326^ experiments. However, the preferential use of arrestins in TORC1-mediated internalization makes this process unique from receptor-mediated endocytosis. Prior studies have shown that a double deletion of *ROG1* and *ROG3* cause significant defects in receptor-mediated Ste2 endocytosis, whereas single deletions of these genes show no significant effect, indicating that these proteins are redundant in their function during mating^47^. However, we find that Rog3 is preferentially used in TORC1-mediated internalization. There are 33 serine and threonine residues in the C-terminal region of Ste2 that can be phosphorylated. Kim et al. found that specific residues contribute to different aspects of both receptor internalization and signaling during mating^85^. Therefore, differential patterns of receptor phosphorylation may lead to preferential recruitment of endocytic adapter proteins, like Rog3, during starvation. Another key difference from ligand-induced internalization is that TORC1-mediated internalization does not seem to require Yck1 or Yck2. It is possible that this stems from the redundant functions of these kinases, with one kinase compensating for the other’s deficiency^48^. However, another kinase, such as Ypk1, may be responsible for direct phosphorylation of the receptor during starvation. Regardless of the enzyme responsible for modifying the receptor, we have found differential use of arrestins that have previously been seen as redundant^47^.

### Activation of TORC2 by inhibition of TORC1

We have found an interesting crossroad in TORC signaling whereby TORC1 signaling appears to directly impact TORC2 activity. TORC1 and TORC2 are stress-responsive complexes that regulate cellular growth and metabolism. These complexes are structurally and functionally distinct. However, there are instances where TORC1 and TORC2 signaling pathways intertwine to tune the magnitude of their responses^86,87^. For example, while TORC1 is the primary regulator of autophagy during nitrogen starvation, TORC2 activity has been suggested to enhance autophagy specifically in the absence of amino acids^86^. We have found that both TORC1 and TORC2 contribute to rapamycin-induced endocytosis of Ste2. This may represent a novel signaling pathway between TORC1 and TORC2 where one regulates the signaling of the other directly at the complex level, but further investigation of this interaction is required.

In this study, we have found that Ypk1 controls receptor endocytosis. Ypk1 is known to play a regulatory role in endocytosis through the TORC2 signaling pathway, but the nature of this regulation is inconclusive^88-91^. Some studies suggest that Ypk1 activity promotes receptor endocytosis^88^, while others suggest that Ypk1 downregulates Rod1 to repress Ste2 internalization^90^. In this study, Ypk1 promotes Ste2 internalization during nutrient deprivation where there is a marked effect on transcriptional output that may be due to the changes in receptor endocytosis.

Our studies provide yet another context where Ypk1 promotes endocytosis, but more work on Ypk1 and endocytosis will need to be done to determine how context impacts their relationship.

Yeast initiate vacuolar targeting of cytoplasmic proteins during the pheromone response, which may represent engagement of selective autophagy such as cytoplasm-to-vacuole transport^62^. One potential explanation is that, as a type of autophagy, it frees metabolites to support growth of the mating projection. However, our data suggests that it serves as a negative regulator of the signaling pathway. We have found that cells use Atg8 to dampen mating pathway output. Atg8 commonly associates with selective autophagy receptors^92-96^. Perhaps mating stimulates autophagic clearance of cytosolic proteins to remove active receptor (and potentially other signaling molecules) from the cytosol. In mammalian cells, GPCR trafficking to various subcellular locations following internalization leads to unique signaling profiles^97-100^. It is possible that Ste2 may likewise signal at the endosome. In this scenario, the cell would rely upon Atg8 and autophagy to suppress endosomal signaling by directing Ste2 to the vacuole.

We found that TOR signaling impacted localization of both mating GPCRs in yeast. There are reports that β_2_-Adrenergic Receptor is targeted to the lysosome in a ligand-dependent manner in mammals^101^. Together, these data suggest that TOR regulation of GPCRs may be broadly conserved. If so, drugs that target TOR in humans may impact the localization or signaling of at least a subset of human GPCRs. GPCRs and the regulation of their signaling is significant to human health and physiology, leading them to being the most targeted molecule by FDA-approved therapeutics^102^. Therefore, understanding signaling cascades that control their output is crucial to therapeutic development for diseases that may arise from dysregulated GPCR signaling, such as heart failure and certain cancers^103-105^. These studies open the doors to avenues of research at the intersection of GPCR and TOR signaling.

## MATERIALS AND METHODS

### Yeast Strains

Strains reported in this study are listed in Table S1. All strains were constructed in the MAT**a** haploid *Saccharomyces cerevisiae* parent strain BY4741, except for those used to visualize Ste3. In which case, strains were constructed in the MATα haploid parent strain BY4742. Strains containing fluorescent labeled, deleted, or truncated genes were created through oligonucleotide-directed homologous recombination, using primers listed in Table S2. Primer amplicons were generated through PCR using Q5 Hot Start High-Fidelity 2X Master Mix (M0494, NEB, United States). G-protein-coupled receptors (Ste2, Ste3, Gpr1) and Ypk1were labeled with a monomeric Envy (mEnvy/Envy^V206K^). EGFP sourced from pFA6a-link-yoEGFP-SpHis5 (Addgene Plasmid #44836)^106^ was used to label and truncate Ste2 simultaneously. Pma1 labeling is discussed below. Genetic deletion mutants and the Avo3 truncation strains were generated by replacing the target locus with a kanamycin resistance cassette (KanMX6) that can be selected for via geneticin treatment. This cassette was sourced from a Yeast Deletion Library (Dharmacon, United States) or from the pFA6a-yo-linkEGFP-KanMX plasmid (Addgene Plasmid #44900)^106^.

Cells were grown in Yeast Extract-Peptone-Dextrose (YPD) growth media at 30°C. PCR products were transformed into parent yeast strains using a standard lithium acetate transformation with single-stranded carrier DNA and PEG^107^. These cultures were grown on standard selective media (SCD his-, SCD leu-, SCD ura-, or YPD G418+) and isolated via auxotrophic or geneticin resistance-based selection. Transformants were verified using PCR and fluorescence microscopy when applicable. PCR verification was performed using One*Taq* 2X Master Mix with Standard Buffer (M0482, NEB, United States) or LongAmp Hot Start *Taq* 2X Master Mix (M0533, NEB, United States).

### Yeast Cell Culture for Microscopy

Cells were grown overnight in 0.22μm-filtered Synthetic Complete + Dextrose (SCD) media at 30°C to an OD600 of 0.2-0.8. To limit cells of nitrogen sources, cells were diluted from SCD media into low nitrogen SCD media (SCD ammonium sulfate-, uracil-, L-histidine-, L-leucine-) and incubated for 4.5 hours prior to imaging. To repress TORC1 activity with rapamycin (0215934691, MP Biomedicals, United States), cells were treated with 0.2µM rapamycin suspended in ethanol for 2 hours prior to imaging. To induce the mating response, cells were treated with 10µM α-factor (RP01002, GenScript, United States) suspended in autoclaved milliQ water for 2 hours prior to imaging.

### Live-cell Widefield Microscopy

All cells were imaged on 2% agarose pads in SCD. Agarose pads were supplemented with the same concentrations of rapamycin or α-factor as in liquid cultures where applicable. Cells were imaged using an inverted widefield fluorescence microscope (IX83, Olympus, Japan) equipped with a Prime 95B CMOS camera (Teledyne Photometrics, United States). All images were acquired using a UPlanXApo 100X/1.45NA Oil objective (N5702400, Olympus, Japan). Z-stack images were collected with a step size of 0.32 µm.

Transmitted light illumination was performed using a 12V 100W halogen light bulb at 5.0V power. Transmitted light exposure time was 200ms. Fluorescence illumination was performed using an X-Cite 120 LEDBoost (Excelitas, United States) at 40% light intensity. EGFP and mEnvy emission was collected using Ex466/40 Em525/50 filter set (GFP-4050B-OFF-ZERO, Semrock, United States) at 3s exposure.

Images were acquired at 200X magnification with an IX3 magnification changer, using the Z before channel setting, with image size being 1200x1200 pixels. The software was cellSens Dimension 2.3 (Olympus, Japan) and images were saved as 16-bit VSI files.

### Analysis of Peripheral GPCR Abundance

Z-stacks of fluorescent and differential interference contrast (DIC) images were acquired as described above. In FIJI, a single slice of the fluorescent and DIC images in the middle of the Z-stack were isolated and saved as 16-bit TIFF files to be used for mask generation and analysis. These images were merged and qualitatively assessed as quality control to ensure that the isolated fluorescent and DIC frame exist in the same focal plane. The saved DIC images were loaded into Cellpose 2.0^108^ for mask generation. In Cellpose 2.0, the cyto2 pretrained model was used to generate masks for individual cells in each image, with a user-defined cell diameter of 80 pixels, a flow threshold (flow_threshold) of 0, and a mask threshold (cellprob_threshold) of 1.5. Any masks which did not encompass the entire cell were fixed or removed. Any masks that overlayed debris, dead cells, or cells that were partially out of frame were removed. Masks were saved as PNG files.

Masks were loaded into MATLAB and eroded with a diamond structuring element (1-pixel radius) to ensure individual cell masks are not touching. Masks were converted into a logical mask, and all connected components with fewer than 5000 pixels were removed. The logical mask was then converted back to a “Double” data format. To convert these into masks that encompass the cell periphery, the masks were first duplicated. The duplicate masks were eroded with a diamond structuring element (7-pixel radius), and the new area encompassed by these masks were removed from the originals. This would leave behind a working mask which covers the edge of each cell. Converting this to a count mask generates the working mask used to measure the fluorescent intensity along the cell edge in individual cells.

Single-slice fluorescent images were loaded into MATLAB. A Gaussian Blur was applied to the image with a standard deviation (sigma) of 50 and subtracted from the original image (performing a pseudo-flatfield background subtraction). The mean fluorescent intensity was calculated from this image within the area defined by the working mask described above for each individual cell.

### Statistical Analysis of Epifluorescence Microscopy Data

In experiments where two sample groups were being compared to each other (Figures 1D and 2E), two-tailed two-sample t-tests of unequal variance were performed. Any p-value produced that was less than 0.05 deemed the two groups to have a difference in means that is statistically significant. In experiments where greater than two groups were being compared (Figures 3B, 4B, 5B, 5D, 6B, 6D), One-Way ANOVAs were performed. Each ANOVA compared four groups: wild-type untreated cells, wild-type cell treated with rapamycin, mutant untreated cells, and mutants treated with rapamycin. The ANOVA results were then processed through a multiple comparison test (Tukey’s honest significance test). As described above, a p-value equal to or less than 0.05 denoted a significant difference between sample groups.

### Mating Gene Transcription Assays

Assays were performed based on the Beta-galactosidase Reporter Gene Assay (Liquid Form) from the Dohlman Lab^70^. Cells were transformed with the plasmid pRS423-pFUS1-LacZ as previously described. Colonies were isolated by growth on standard selective media (SCD his-). Cultures were grown in SCD his-media at 30°C to an OD600 of 0.1-0.8. Cells were pelleted at 2500rpm for 5min and resuspended in SCD his-media to obtain an OD_600_ of 0.6-0.8. In a polystyrene 96-well plate, 90µL of cells were added to each well per row. Each row of wells containing cells represented an individual biological replicate, as these cells were grown from different transformant colonies. Then, 10µL of α-factor was added to each well, whereby each column of wells contained different concentrations of the α-factor: e.g. 0µM, 0.003µM, 0.01µM, 0.03µM, 0.1µM… 300µM. Cells were gently shaken and incubated at 30°C for 90min. Solutions of 1mM fluorescein di-β-galactopyranoside (F1179, Invitrogen) diluted in 25mM potassium phosphate buffer pH 7.2 and 5% Triton X-100 diluted in 250mM potassium phosphate buffer pH 7.2 were mixed in equal parts. After the 90min incubation, 20µL of this solution was added to each well, cells were gently shaken, covered in aluminum foil, and incubated at 37°C for 90min. The reaction was quenched with 20µL of 1M sodium bicarbonate and read on a fluorescent plate reader (BioTek Synergy 2, Agilent, United States) at an excitation wavelength of 485/20 and emission wavelength of 528/20. Fluorescent intensity values were normalized to cell culture density and averages were fit to a Hill Equation using the Curve Fitting Toolbox 3.8 in MATLAB to determine the E_max_, EC_50_, and Hill slope of the generated dose response curve with their respective 95% confidence intervals. Differences in these values were considered statistically significant if their confidence intervals did not overlap. Lower bounds of these values were set to zero and upper bounds remained infinite.

### Quantitative Mating Assays

Assays were performed based on the Quantitative Mating Assays described by Sprague Jr et al^71^. Cultures of MAT**a** (BY4741) and MATα (BY4742) cells were grown in YPD at 30°C overnight to an OD600 of 0.2-0.8. Cell densities of these cultures in cells/mL were then determined using a hemocytometer. Then, 10^7^ MATα cells were mixed with 2x10^6^ MAT**a** cells. This suspension was then filtered through a 0.45µm-pore 25mm-diameter nitrocellulose filter disk, such that the liquid media was discarded, leaving the cells to lay on the filter. The filter was then transferred to a YPD plate (YPD rapamycin+ if cells were pretreated with 0.2µM rapamycin) and incubated at 30°C for 5h. Cells were then resuspended in 2mL of SCD media and further diluted such that a countable number of colonies was achieved once plated. Cells were plated on YPD and incubated at 30°C until colonies were visible (1 day for untreated cells, 2 days for rapamycin-treated cells). Cells were then replica plated onto SCD lys- and SCD lys- met- cys- to select for MAT**a** + diploid cells and diploid cells respectively. These plates were incubated 30°C for 1 day, and colonies from each plate were counted. Mating efficiency was determined by calculating the percentage of cells that mated (SCD lys- met- cys- colonies) out of all mating competent cells in the mating suspension (SCD lys-colonies). To determine whether the mean mating efficiencies of rapamycin-treated and untreated groups were significantly different, we performed a 1-tailed Two-Sample t-test of unequal variance. A p-value of less than 0.05 was considered statistically significant.

### Live-cell Confocal Microscopy

All cells were imaged on 2% agarose pads in SCD. Agarose pads were supplemented with 10µM α-factor, the same concentration as in liquid cultures. Images were acquired using a point scanning confocal unit (LSM 980, Carl Zeiss Microscopy, Germany) on a Zeiss Axio Observer 7 inverted microscope (409000-9663-000, Carl Zeiss Microscopy, Germany) equipped with a C Plan-Apochromat 63X/1.4NA Oil objective lens (421782-9900-799, Carl Zeiss Microscopy, Germany). Samples were placed into an Incubator XLmulti S2 DARK (ref: 411857-9310-000, Carl Zeiss Microscopy, Germany), warmed at 30°C using the Temp Module S1 (ref:411860-9010-000, Carl Zeiss Microscopy, Germany) and the heating unit XL S2 (ref:411857-9031-000, Carl Zeiss Microscopy, Germany). Z-stack images were collected with a step size of 0.17 µm with the Motorized Scanning Stage 130x100 STEP Set LSM (ref: 409000-9420-000, Carl Zeiss Microscopy, Germany) mounted on the universal mounting frame K-M (ref: 432341-9100-000, Carl Zeiss Microscopy, Germany) and the insert mounting frame K-M for specimen slides (ref: 432340-9040-000, Carl Zeiss Microscopy, Germany).

mEnvy fluorescence was excited with a 488nm diode laser at 3% power and collected using an Airyscan 2 detector at 800V with 495-555nm BP and 660nm LP filters through a pinhole at 5.25AU. ytdTomato fluorescence was excited with a 561nm diode laser at 3% power and collected using an Airyscan 2 detector at 850V with 420-480nm BP and 570-630 BP filters through a pinhole at 5.00AU. Airyscan images were processed manually in 3D with a strength of the deconvolution set to 7.5.

Images were frame-scanned sequentially in SR mode, with a pixel time of 0.66μs, a frame time of 486.6μs, a zoom factor of 4X, using bidirectional scanning. Images were acquired such that each track were acquired before moving to the next Z-step. Images were 768x768 pixels, each pixel being 0.043x0.043μm. The software was Zen Blue 3.8 (Carl Zeiss Microscopy, Germany) and images were saved as 16-bit CZI files.

### Analysis of GPCR Abundance at the Vacuolar Membrane and Lumen

Using FIJI, background was subtracted from GFP and RFP images with a rolling ball radius of 50 pixels. Linescans were drawn through the largest vacuolar body in each cell. The mean fluorescent intensities of Vph1-Tomato (vacuolar membrane) and Ste2-mEnvy (mating receptor) were measured at each position across the linescans. Using MATLAB, the two local maxima of Vph1-Tomato fluorescence and their immediately flanking positions were labeled as the vacuolar membrane. At these positions, the mean fluorescent intensities of Ste2-mEnvy were averaged to determine receptor levels at the vacuolar membrane. Between these positions, Ste2-mEnvy intensities were averaged to determine receptor levels in the vacuolar lumen. For each cell, Ste2 levels at the vacuolar membrane were divided by receptor levels in the lumen. Cells were then classified as expressing Phenotype 1 (full vacuoles) if their lumen:membrane ratios were greater than 1. Cells were classified as expressing Phenotype 2 (empty vacuoles) if their lumen:membrane ratios were less than 1. Violin plots were generated in MATLAB using Violinplot-Matlab^109^.

### Mutagenesis of GFP Envy in an Epitope-Tagging Plasmid

The pFA6a-link-GFPEnvy-SpHis5 epitope tagging plasmid (Addgene Plasmid #60782)^110^ was mutagenized such that the expressed Envy protein was incapable of dimerization. Val206 of Envy was mutagenized to Lys with the Q5 Site-Directed Mutagenesis Kit (E0554S, NEB, United States) using primers listed in Table S2. The mutated product was transformed into chemically competent *Escherichia coli* through a High Efficiency Transformation (C2987H, NEB, United States). Colonies were isolated via growth on selective media (LB carbenicillin+). Colonies were grown in liquid selective media, and plasmids were purified using a Monarch Plasmid Miniprep Kit (T1010, NEB, United States). Successful mutagenesis was verified through DNA sequencing (Plasmidsaurus).

### Cloning pRSII405-*PMA1-mRUBY2*

A plasmid construct containing *PMA1-mRUBY2* was generated in a pRSII405 vector via Gibson Assembly. The sequence for *PMA1* was amplified from BY4741 genomic DNA, and the *mRUBY2* sequence was amplified from pFA6a-link-yomRuby2-Kan (Addgene Plasmid #44953)^106^. Both amplicons were generated through PCR with Q5 Hot Start High-Fidelity 2X Master Mix (M0494, NEB, United States) using primers listed in Table S2. The vector, pRSII415 (Addgene Plasmid #35440)^111^ was digested by SacII (R1057, NEB, United States) and HindIII-HF (R3104, NEB, United States). The cut vector was verified via band separation using gel electrophoresis, and the vector was extracted from the gel using a Monarch DNA Gel Extraction Kit (T1020, NEB, United States). Gibson Assembly of the three generated fragments was performed using NEBuilder HiFi DNA Assembly Master Mix (E2621, NEB, United States). The resulting assembly mixture was transformed into chemically competent *Escherichia coli* through a High Efficiency Transformation (C2987H, NEB, United States). Colonies were isolated via growth on selective media (LB carbenicillin+). Colonies were grown in liquid selective media, and plasmids were purified using a Monarch Plasmid Miniprep Kit (T1010, NEB, United States). Constructs were then verified by restriction digest using SacI-HF (R3156, NEB, United States) and SalI-HF (R3138, NEB, United States) and through DNA sequencing (Plasmidsaurus, United States). Verified plasmids were digested BamHI-HF (R3136, NEB, United States) and transformed into yeast using the transformation methods described above. Colonies were grown on standard selective media (SCD leu-) and isolated via auxotrophic selection. Transformants were verified using PCR and fluorescence microscopy. PCR verification was performed using One*Taq* 2X Master Mix with Standard Buffer (M0482, NEB, United States).

## Supplementary Materials

Fig. S1. Single genetic deletions that do not block rapamycin-induced endocytosis.

Fig. S2. Plasma membrane intensity data without normalization.

Table S1. Yeast Strains.

Table S2. Primer List.

## Acknowledgments

The authors wish to thank Claire Gordy for helpful discussions. The authors would like to acknowledge the University of Maine Microscopy and Image Analysis Core Facility (RRID:SCR_025784) for use of Zeiss LSM980 Confocal Microscope and Zhengxin Ma for helpful training and discussion. The MIAC facility was supported by Award number 1P20GM144265 from the National Institutes of Health.

## Author contributions

Conceptualization: NRL, JBK

Data Curation: NRL

Formal Analysis: NRL

Funding Acquisition: JBK, TD

Investigation: NRL, TD, SS, CJ

Methodology: NRL, JBK

Project Administration: NRL, JBK

Resources: JBK

Software: NRL, JBK

Supervision: JBK

Validation: NRL, JBK

Visualization: NRL, JBK

Writing – Original Draft: NRL, JBK

Writing – Review & Editing: NRL, JBK

## Data and materials availability

MATLAB scripts used for the analysis of receptor localization at the plasma membrane are available on GitHub (https://github.com/Kelley-Lab-Computational-Biology).

