## Supplemental FIgures for "TOR signaling regulates GPCR levels on the plasma membrane and suppresses the *Saccharomyces cerevisiae* mating pathway"

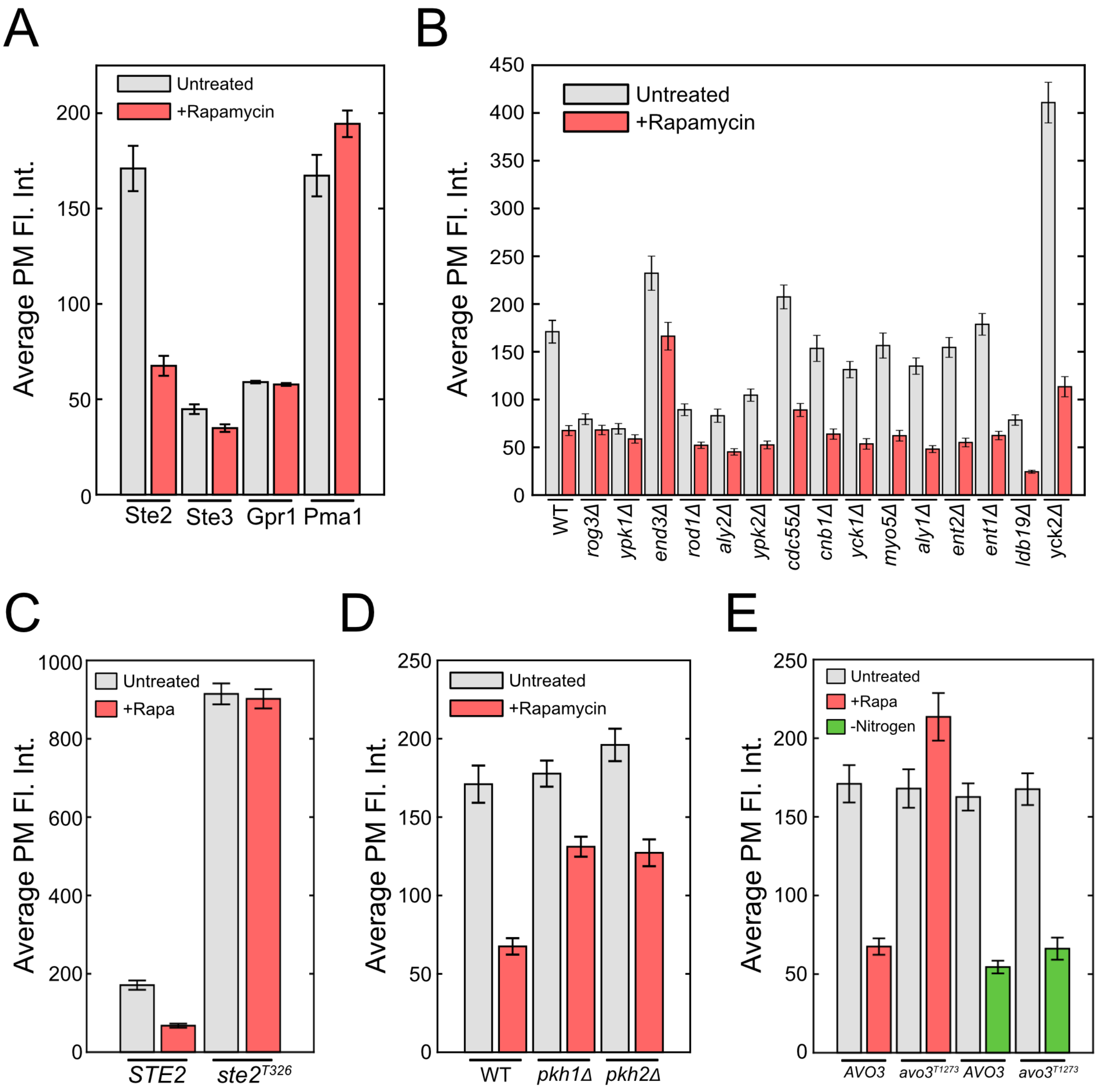


**Figure S1. Plasma membrane intensity data without normalization.** (**A**) Unnormalized data reported in Figure 2. (**B**) Unnormalized data reported in Figure 3. (**C**) Unnormalized data reported in Figure 4. (**D**) Unnormalized data reported in Figure 5. (**E**) Unnormalized data reported in Figure 6.

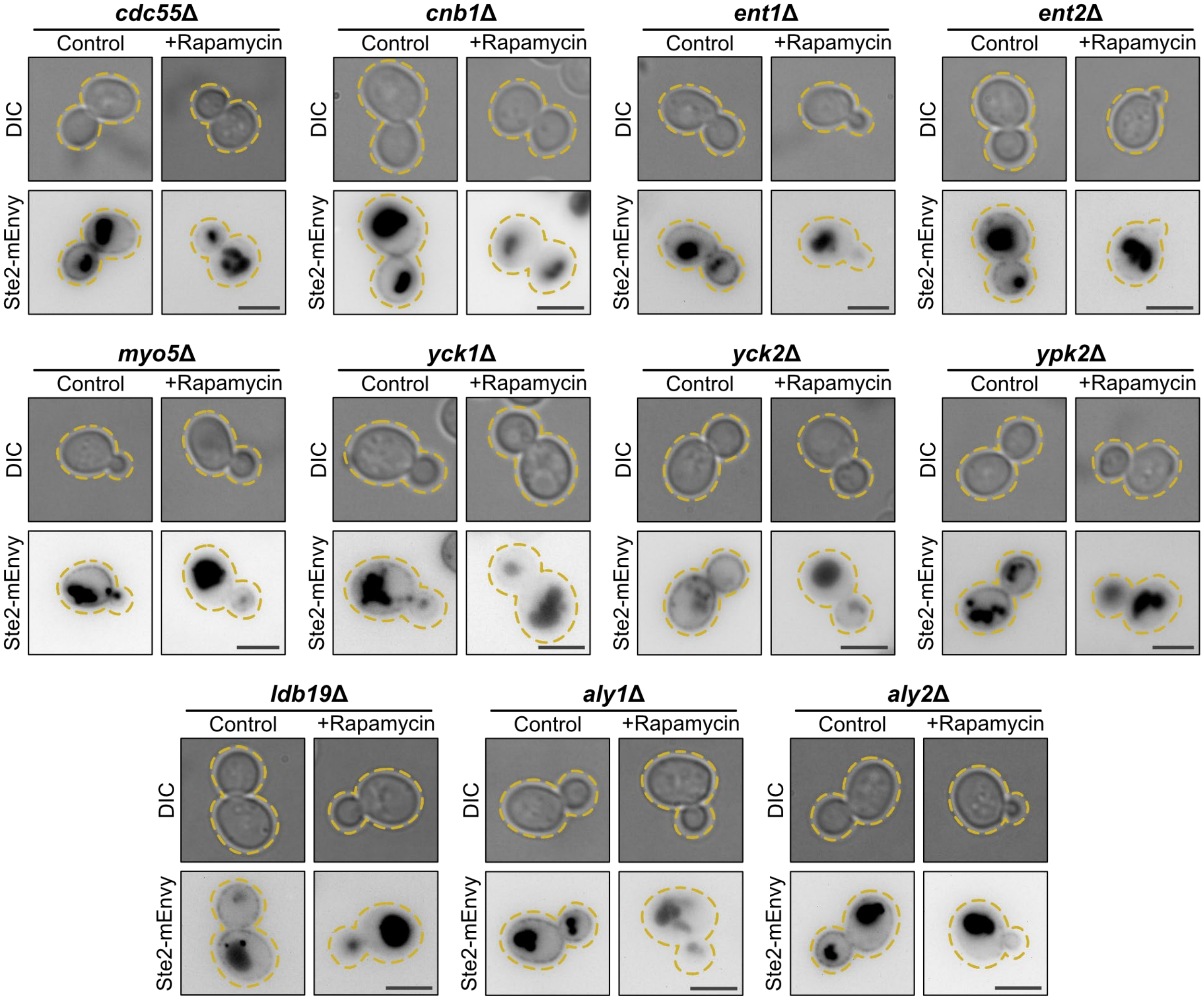


**Figure S2. Single genetic deletions that do not block rapamycin-induced endocytosis.** Representative images of genetic deletion mutants that do not block Ste2 internalization in the presence of 0.2µM rapamycin. Scale bars represent 4µm.

**Table S1. Strain List.**

| ***Strain*** | ***Parent*** | ***Description*** |
|---|---|---|
| BY4741 | S288C | *his3Δ1*<br>*leu2Δ0*<br>*met15Δ0*<br>*ura3Δ0* |
| BY4742 | S288C | *his3Δ1*<br>*leu2Δ0*<br>*lys2Δ0*<br>*ura3Δ0* |
| *STE2-mEnvy* | BY4741 | *STE2:Envy*$^{V206K}$*::HIS3* |
| *STE3-mEnvy* | BY4742 | *STE3:Envy*$^{V206K}$*::HIS3* |
| *GPR1-mEnvy* | BY4741 | *GPR1:Envy*$^{V206K}$*::HIS3* |
| *PMA1-mRuby2* | BY4741 | *PMA1:mRuby2::LEU2* |
| *STE2-mEnvy aly1Δ* | BY4741 | *STE2:Envy*$^{V206K}$*::HIS3*<br>*aly1Δ::KanMX6* |
| *STE2-mEnvy aly2Δ* | BY4741 | *STE2:Envy*$^{V206K}$*::HIS3*<br>*aly2Δ::KanMX6* |
| *STE2-mEnvy cdc55Δ* | BY4741 | *STE2:Envy*$^{V206K}$*::HIS3*<br>*cdc55Δ::KanMX6* |
| *STE2-mEnvy cnb1Δ* | BY4741 | *STE2:Envy*$^{V206K}$*::HIS3*<br>*cnb1Δ::KanMX6* |
| *STE2-mEnvy end3Δ* | BY4741 | *STE2:Envy*$^{V206K}$*::HIS3*<br>*end3Δ::KanMX6* |
| *STE2-mEnvy ent1Δ* | BY4741 | *STE2:Envy*$^{V206K}$*::HIS3*<br>*ent1Δ::KanMX6* |
| *STE2-mEnvy ent2Δ* | BY4741 | *STE2:Envy*$^{V206K}$*::HIS3*<br>*ent2Δ::KanMX6* |
| *STE2-mEnvy ldb19Δ* | BY4741 | *STE2:Envy*$^{V206K}$*::HIS3*<br>*ldb19Δ::KanMX6* |
| *STE2-mEnvy myo5Δ* | BY4741 | *STE2:Envy*$^{V206K}$*::HIS3*<br>*myo5Δ::KanMX6* |

| | | |
|---|---|---|
| *STE2-mEnvy rod1Δ* | BY4741 | *STE2:Envy*$^{V206K}$*::HIS3 rod1Δ::KanMX6* |
| *STE2-mEnvy rog3Δ* | BY4741 | *STE2:Envy*$^{V206K}$*::HIS3 rog3Δ::KanMX6* |
| *STE2-mEnvy yck1Δ* | BY4741 | *STE2:Envy*$^{V206K}$*::HIS3 yck1Δ::KanMX6* |
| *STE2-mEnvy yck2Δ* | BY4741 | *STE2:Envy*$^{V206K}$*::HIS3 yck2Δ::KanMX6* |
| *STE2-mEnvy ypk1Δ* | BY4741 | *STE3:Envy*$^{V206K}$*::HIS3 ypk1Δ::KanMX6* |
| *STE2-mEnvy ypk2Δ* | BY4741 | *STE2:Envy*$^{V206K}$*::HIS3 ypk2Δ::KanMX6* |
| *ste2*$^{T326}$*-EGFP* | BY4741 | *ste2*$^{T326}$*:EGFP::HIS3* |
| *ypk1Δ* | BY4741 | *ypk1Δ::KanMX6* |
| *STE2-mEnvy avo3*$^{T1273}$ | BY4741 | *STE2:Envy*$^{V206K}$*::HIS3 avo3*$^{T1273}$*::KanMX6* |
| *STE2-mEnvy pkh1Δ* | BY4741 | *STE2:Envy*$^{V206K}$*::HIS3 pkh1Δ::KanMX6* |
| *STE2-mEnvy pkh2Δ* | BY4741 | *STE2:Envy*$^{V206K}$*::HIS3 pkh2Δ::KanMX6* |
| *STE2-mEnvy VPH1-Tomato* | BY4741 | *STE2:Envy*$^{V206K}$*::HIS3 VPH1:ytdTomato::URA3* |
| *STE2-mEnvy VPH1-Tomato atg8Δ* | BY4741 | *STE2:Envy*$^{V206K}$*::HIS3 VPH1:ytdTomato::URA3 atg8Δ::KanMX6* |
| *atg8Δ* | BY4741 | *atg8Δ::KanMX6* |

**Table S2. Primer List.**

| *Primer Name* | *Sequence* | *Description* |
|---|---|---|
| *WSM-7* | 5' GGAAGCCAGAAAGTTCTGGACTGAAGATAATAATAATTTAGGTGACGGTGCTGGTTTA 3' | Amplify fluorescent label to tag Ste2 |
| *WSM-8* | 5' GAAGGTCACGAAATTACTTTTTCAAAGCCGTAAATTTTGATCGATGAATTCGAGCTCG 3' | Amplify fluorescent label to tag Ste2 |
| *WSM-11* | 5' GATGCTAAAAGCAGTCTCAG 3' | Verify Ste2 labeling |
| *WSM-12* | 5' GAGAGTTCTAGATCATGGCA 3' | Verify Ste2 labeling |
| *NLM-1* | 5' CGAGGTCGACGGTATCGATACCAAAGGCTAAGGACGCTTTG 3' | Amplify *PMA1* ORF for Gibson Assembly |
| *NLM-2* | 5' CACCGTCACCGGTTTCCTTTTCGTGTTGAGTAG 3' | Amplify *PMA1* ORF for Gibson Assembly |
| *NLM-3* | 5' AAAGGAAACCGGTGACGGTGCTGGTTTAATTAAC 3' | Amplify *mRUBY2* ORF for Gibson Assembly |
| *NLM-4* | 5' AAAAGCTGGAGCTCCACCGCTTACTTATACAATTCATCCATACCACC 3' | Amplify *mRUBY2* ORF for Gibson Assembly |
| *NLM-5* | 5' AAGCTCAAACGAACATAGTTCAGAAAATACTGCAGGCCCTGGTGACGGTGCTGGTTTA 3' | Amplify fluorescent label to tag Ste3 |
| *NLM-6* | 5' AATACTCCTAGTCCAGTAAATATAATGCGACACTCTTGTGTCGATGAATTCGAGCTCG 3' | Amplify fluorescent label to tag Ste3 |
| *NLM-7* | 5' CGGCATAGATTTGATAGCCTTCTTAAGAAATGGACCATTAGGTGACGGTGCTGGTTTA 3' | Amplify fluorescent label to tag Gpr1 |
| *NLM-8* | 5' TTCCTTACTTTCCATTTTCAAACATCGCGATACAAAAACTTCGATGAATTCGAGCTCG 3' | Amplify fluorescent label to tag Gpr1 |
| *NLM-9* | 5' ACACCAATATCACAAGCGCA 3' | Verify Ste3 labeling |
| *NLM-10* | 5' TCTGCTAATCGACTTTTGGAGC 3' | Verify Ste3 labeling |
| *NLM-11* | 5' TCTACCCGGGTTGAAATTTGC 3' | Verify Gpr1 labeling |

| | | |
|---|---|---|
| *NLM-12* | 5' ACGAGCACTCATCCATTTTCA 3' | Verify Gpr1 labeling |
| *NLM-15* | 5' AGAACCGACCAAGACGTGTT 3' | Amplify deletion cassette at *ATG8* ORF from Yeast Deletion Library |
| *NLM-16* | 5' GTACGTTAAGAACAGCGGCA 3' | Amplify deletion cassette at *ATG8* ORF from Yeast Deletion Library |
| *NLM-35* | 5' TCCCTCGTTCACAGAAAGTCT 3' | Verify Pma1 labeling |
| *NLM-36* | 5' TGCATCACAGGTCCGTTAGA 3' | Verify Pma1 labeling |
| *NLM-63* | 5' CCTTTTGATGTTACCCCGCC 3' | Amplify deletion cassette at *YCK1* ORF from Yeast Deletion Library |
| *NLM-64* | 5' AAAGGGGCAAAGGTGTGAAG 3' | Amplify deletion cassette at *YCK1* ORF from Yeast Deletion Library |
| *NLM-65* | 5' TCCACGTAGAACATCGCAGT 3' | Verify *YCK1* deletion |
| *NLM-66* | 5' ACAGGAATCGAATGCAACCG 3' | Verify *YCK1* deletion |
| *NLM-67* | 5' TCATTAAAGTGTGGGCTGTGG 3' | Amplify deletion cassette at *YCK2* ORF from Yeast Deletion Library |
| *NLM-68* | 5' TGCAAATTGAAAGAGGGTAAACA 3' | Amplify deletion cassette at *YCK2* ORF from Yeast Deletion Library |
| *NLM-69* | 5' GTCTTGATGCTCTGAAGGCG 3' | Verify *YCK2* deletion |
| *NLM-70* | 5' TCCGACTCGTCCAACATCAA 3' | Verify *YCK2* deletion |
| *NLM-71* | 5' ACCAACAGTCCGCACATAGA 3' | Verify *YPK1* deletion |

| | | |
|---|---|---|
| *NLM-72* | 5’ CCATGAGACACAAGCCACAC 3’ | Verify *YPK1* deletion |
| *NLM-79* | 5’ AGTTCCTGTTTTGCCAATGGT 3’ | Amplify deletion cassette at *MYO5* ORF from Yeast Deletion Library |
| *NLM-80* | 5’ GGAATTACCGACGCTCCATT 3’ | Amplify deletion cassette at *MYO5* ORF from Yeast Deletion Library |
| *NLM-81* | 5’ TTGAGTTCTGCCGTTCAAGC 3’ | Verify *MYO5* deletion |
| *NLM-82* | 5’ TAACGGCTCCAAGATTGTGC 3’ | Verify *MYO5* deletion |
| *NLM-83* | 5’ TGGGTCAAATTATCGCGTATACAAATATACATA TAGTAACGACATGGAGGCCCAGAATAC 3’ | Amplify deletion cassette with homology to *YPK2* UTRs |
| *NLM-84* | 5’ AAATTCCGTCCGGCTCGGCTCGGCTTGCTTCG GCTTGCTTCAGTATAGCGACCAGCATTC 3’ | Amplify deletion cassette with homology to *YPK2* UTRs |
| *NLM-85* | 5’ TGGCGTGGTTGAACATCTTG 3’ | Verify *YPK2* deletion |
| *NLM-86* | 5’ TCCGACTCGTCCAACATCAA 3’ | Verify *YPK2* deletion (with NLM-85), *ROD1* deletion (with RAM-15), and *ROG3* deletion (with RAM-17) |
| *NLM-89* | 5’ CCGATGAGGCAAAATATGGTGT 3’ | Verify *ATG8* deletion |
| *NLM-90* | 5’ CGAACTCTTTCCCATTGTCTGT 3’ | Verify *ATG8* deletion |
| *NLM-91* | 5’ GAGGGGTCAGAAGATGCAGA 3’ | Amplify deletion cassette at *ALY1* ORF from Yeast Deletion Library |
| *NLM-92* | 5’ CCGGTACTTTTCCCAGACGA 3’ | Amplify deletion cassette at *ALY1* ORF from Yeast Deletion Library |

| | | |
|---|---|---|
| *NLM-93* | 5’ ATTCTGGGGAGGAGCAAGTC 3’ | Verify *ALY1* deletion |
| *NLM-94* | 5’ CCAACCCGAGGAGAAATTGC 3’ | Verify *ALY1* deletion |
| *NLM-95* | 5’ ACGCCTTCACCTATCACTCT 3’ | Amplify deletion cassette at *ALY2* ORF from Yeast Deletion Library |
| *NLM-96* | 5’ GCGGGAAGAAGTCAAAAGACA 3’ | Amplify deletion cassette at *ALY2* ORF from Yeast Deletion Library |
| *NLM-97* | 5’ GATGTCGAACGGAGAGCAAC 3’ | Verify *ALY2* deletion |
| *NLM-98* | 5’ TTGGAACGGCTGAAGAAACG 3’ | Verify *ALY2* deletion |
| *NLM-99* | 5’ CACATCAAGACCACTGCGAG 3’ | Amplify deletion cassette at *CNB1* ORF from Yeast Deletion Library |
| *NLM-100* | 5’ AGATGGTCTGTCTCCTAGCA 3’ | Amplify deletion cassette at *CNB1* ORF from Yeast Deletion Library |
| *NLM-101* | 5’ GGGAAATGGGTTGTGGACTT 3’ | Verify *CNB1* deletion |
| *NLM-102* | 5’ AATCAGCGGGTTTCCTCCTT 3’ | Verify *CNB1* deletion |
| *NLM-107* | 5’ AGCTCCACCTCAAAGACCAA 3’ | Verify *CDC55* deletion |
| *NLM-108* | 5’ TCCGACTCGTCCAACATCAA 3’ | Verify *CDC55* deletion |
| *NLM-111* | 5’ GGAGAGATCTTACGCATAAAGAAATATAATATAG CGCACAGACATGGAGGCCCAGAATAC 3’ | Amplify deletion cassette with homology to *CDC55* UTRs |
| *NLM-112* | 5’ GGGATAAAAAAAAGTAAGGGAAAATAAGGAATT ATTATAACAGTATAGCGACCAGCATTC 3’ | Amplify deletion cassette with homology to *CDC55* UTRs |

| | | |
|---|---|---|
| *NLM-115* | 5’ GTTCCTCTCATATCAACAAACATTAATACAGTTCCTGAAAGACATGGAGGCCCAGAATAC 3’ | Amplify deletion cassette with homology to *YPK1* UTRs |
| *NLM-116* | 5’ TATGTCATGAGTAAACTAGTTGATAATGTATTCACTAAGTCAGTATAGCGACCAGCATTC 3’ | Amplify deletion cassette with homology to *YPK1* UTRs |
| *NLM-117* | 5’ TACAATTTCTTTGTATACAGCGGGAAATTGACACTTCAAAGTTCCTCTCATATCAACAAA 3’ | Extend homology of NLM-115/NLM-116 amplicon to *YPK1* UTR. |
| *NLM-118* | 5’ AGTAAGTAACGGAAAAGAAAACCTTTCTTTTTATATAAAGTATGTCATGAGTAAACTAGT 3’ | Extend homology of NLM-115/NLM-116 amplicon to *YPK1* UTR. |
| *NLM-150* | 5’ AACAGAAGGGTGCCAACATG 3’ | Verify *PKH1* deletion |
| *NLM-151* | 5’ CCTGAAATACATGCCCGAAGA 3’ | Verify *PKH1* deletion |
| *NLM-154* | 5’ GCGAACATTTCCCGATCCTT 3’ | Amplify deletion cassette at *PKH2* ORF from Yeast Deletion Library |
| *NLM-155* | 5’ AAGCGTTGCCTTTGTGAGC 3’ | Amplify deletion cassette at *PKH2* ORF from Yeast Deletion Library |
| *NLM-156* | 5’ AGCTATTGACGAAGGCCCAT 3’ | Verify *PKH2* deletion |
| *NLM-157* | 5’ GCAGTTAATCCAAGGTGCCA 3’ | Verify *PKH2* deletion |
| *NLM-162* | 5’ AGCGGATGGTCGTTCTTCTC 3’ | Verify *END3* deletion |
| *NLM-163* | 5’ GCTGAAGCGTGAGAATGAGT 3’ | Verify *END3* deletion |
| *NLM-166* | 5’ AAGCGATGGCGACTTGATAGAAAAATGCTTACCCATCTAGGACATGGAGGCCCAGAATAC 3’ | Amplify deletion cassette to truncate *AVO3* |
| *NLM-167* | 5’ TTGTGACTATATACATTTATACATGCGGCCCTTTTTTGCTCAGTATAGCGACCAGCATTC 3’ | Amplify deletion cassette to truncate *AVO3* |

| | | |
|---|---|---|
| *NLM-168* | 5’ AGCCTTAGAAACCGACACGA 3’ | Verify *AVO3* truncation |
| *NLM-169* | 5’ TCTGCTGAAACGGAACTCCC 3’ | Verify *AVO3* truncation |
| *NLM-187* | 5’ GCACGTGTACTTGCTTGAATACTGCTACTA TATCATTAATGACATGGAGGCCCAGAATAC 3’ | Amplify deletion cassette with homology to *PKH1* UTRs |
| *NLM-188* | 5’ TGTCTTACATATGCATATATATATATTATCAA GCACAGTTCAGTATAGCGACCAGCATTC 3’ | Amplify deletion cassette with homology to *PKH1* UTRs |
| *NLM-189* | 5’ TATTGGAAAGGCCGGTAAAGATAACAGGGATCTC TGAAAAGACATGGAGGCCCAGAATAC 3’ | Amplify deletion cassette with homology to *END3* UTRs |
| *NLM-190* | 5’ AAATATTACACATTCATGTACATAAAATTAATTATC GGTGCAGTATAGCGACCAGCATTC 3’ | Amplify deletion cassette with homology to *END3* UTRs |
| *NLM-193* | 5’ TATAATCAGACGATTGTAATATAACCATAGAGTTAG TGGGTATTGGAAAGGCCGGTAAAG 3’ | Extend homology of NLM-189/NLM-190 amplicon to *END3* UTR. |
| *NLM-194* | 5’ TTCAAATCAAAAAAGTTTACAAGTGAAATAACAAA CAGTAAATATTACACATTCATGTAC 3’ | Extend homology of NLM-189/NLM-190 amplicon to *END3* UTR. |
| *NLM-242* | 5’ GGAAGTCGCTGTTGCTAGTGCAAGCTCT TCCGCTTCAAGCGGTGACGGTGCTGGTTTA 3’ | Amplify fluorescent label to tag Vph1 |
| *NLM-243* | 5’ AGTACTTAAATGTTTCGCTTTTTTTAAAAG TCCTCAAAATTCGATGAATTCGAGCTCG 3’ | Amplify fluorescent label to tag Vph1 |
| *NLM-244* | 5’ CCCTCATCAAGCAAAGGTCC 3’ | Verify Vph1 labeling |
| *NLM-245* | 5’ GAACACCATCAACGGATCGA 3’ | Verify Vph1 labeling |
| *RAM-1* | 5’ CCCTACCCGACTGCAAACTA 3’ | Amplify deletion cassette at *ROD1* ORF from Yeast Deletion Library |
| *RAM-2* | 5’ TGTTGAGGAAGAAGTGCCAAG 3’ | Amplify deletion cassette at *ROD1* ORF |

| | | |
|---|---|---|
| | | from Yeast Deletion Library |
| *RAM-5* | 5’ GTTATGGACCCGGAGAGAGG 3’ | Amplify deletion cassette at *ROG3* ORF from Yeast Deletion Library |
| *RAM-6* | 5’ GTGTCGCAGTCCATAGAAGG 3’ | Amplify deletion cassette at *ROG3* ORF from Yeast Deletion Library |
| *RAM-9* | 5’ GGCCAGGTTGCTAGTGACTA 3’ | Amplify deletion cassette at *LDB19* ORF from Yeast Deletion Library |
| *RAM-10* | 5’ TGCCCACCCTTTATTTTGCC 3’ | Amplify deletion cassette at *LDB19* ORF from Yeast Deletion Library |
| *RAM-13* | 5’ ACCGTTGATGCTGATGAGGA 3’ | Verify *LDB19* deletion |
| *RAM-14* | 5’ CCCTCAAGCATCGCAGTTT 3’ | Verify *LDB19* deletion |
| *RAM-15* | 5’ CAAATCACTACAAGCCCGCA 3’ | Verify *ROD1* deletion (with NLM-86) |
| *RAM-17* | 5’ AGTATCGATGCGCTGAGTGA 3’ | Verify *ROG3* deletion (with NLM-86) |
| *SLM-3* | 5' TAGGCTTCTTACCGGCAGAG 3' | Amplify deletion cassette at *ENT1* ORF from Yeast Deletion Library |
| *SLM-4* | 5' AGGACATGAGTACAGAGCACA 3' | Amplify deletion cassette at *ENT1* ORF from Yeast Deletion Library |
| *SLM-5* | 5' AGATAGATGGCCGAACGTGG 3' | Amplify deletion cassette at *ENT2* ORF from Yeast Deletion Library |
| *SLM-6* | 5' GACTCCAAAGGTGAATCTGGC3' | Amplify deletion cassette at *ENT2* ORF from Yeast Deletion Library |

| | | |
|---|---|---|
| *SLM-7* | 5' GGGAGAAGGAGTACCTTCTG 3' | Verify *ENT1* deletion |
| *SLM-8* | 5' CCAGAATTATCTTTCGGGCC 3' | Verify *ENT1* deletion |
| *SLM-10* | 5' CGAAGATGGCCGAATGATTGG 3' | Verify *ENT2* deletion |
| *SLM-21* | 5’ TGTGGGCATCTCTGGTTTGA 3’ | Verify *ENT2* deletion |
| *CJM-1* | 5’ ACAACATCCACAGATAGGTTTTATCCAGGC ACGCTGTCTAGCGGTGACGGTGCTGGTTTA 3’ | Amplify fluorescent label to tag and truncate Ste2 |
| *CJM-2* | 5’ CGAAGGTCACGAAATTACTTTTTCAAAGC CGTAAATTTTGACGATGAATTCGAGCTCGTT 3’ | Amplify fluorescent label to tag and truncate Ste2 |
| *CJM-3* | 5’ CAATGTGGGCCACGGCTGCTAA 3’ | Verify labeling and truncation of Ste2 |
| *CJM-4* | 5’ ACAGCGTACCTTTAGACACGTGGG 3’ | Verify labeling and truncation of Ste2 |
| *JKM-70* | 5’ AACCCAATCTaagCTATCTAAAGACC 3’ | Mutate Envy $Val^{206}$ to Lys |
| *JKM-71* | 5’ GATAGGTAGTGGTTGTCTG 3’ | Mutate Envy $Val^{206}$ to Lys |
